# Genomic and epidemiological monitoring of yellow fever virus transmission potential

**DOI:** 10.1101/299842

**Authors:** Faria N. R., Kraemer M. U. G., Hill S. C., Goes de Jesus J., de Aguiar R. S., Iani F. C. M., Xavier J., Quick J., du Plessis L., Dellicour S., Thézé J., Carvalho R. D. O., Baele G., Wu C.-H., Silveira P. P., Arruda M. B., Pereira M. A., Pereira G. C., Lourenço J., Obolski U., Abade L., Vasylyeva T. I., Giovanetti M., Yi D., Weiss D.J., Wint G. R. W., Shearer F. M., Funk S., Nikolai B., Adelino T. E. R., Oliveira M. A. A., Silva M. V. F., Sacchetto L., Figueiredo P. O., Rezende I. M., Mello E. M., Said R. F. C., Santos D. A., Ferraz M. L., Brito M. G., Santana L. F., Menezes M. T., Brindeiro R. M., Tanuri A., dos Santos F. C. P., Cunha M. S., Nogueira J. S., Rocco I. M., da Costa A. C., Komninakis S. C. V., Azevedo V., Chieppe A. O., Araujo E. S. M., Mendonça M. C. L., dos Santos C. C., dos Santos C. D., Mares-Guia A. M., Nogueira R. M. R., Sequeira P. C., Abreu R. G., Garcia M. H. O., Alves R. V., Abreu A. L., Okumoto O., Kroon E. G., de Albuquerque C. F. C., Lewandowski K., Pullan S. T., Carroll M., Sabino E. C., Souza R. P., Suchard M. A., Lemey P., Trindade G. S., Drumond B. P., Filippis A. M. B., Loman N. J., Cauchemez S., Alcantara L. C. J., Pybus O. G.

## Abstract

The yellow fever virus (YFV) epidemic that began in Dec 2016 in Brazil is the largest in decades. The recent discovery of YFV in Brazilian *Aedes sp.* vectors highlights the urgent need to monitor the risk of re-establishment of domestic YFV transmission in the Americas. We use a suite of epidemiological, spatial and genomic approaches to characterize YFV transmission. We show that the age- and sex-distribution of human cases in Brazil is characteristic of sylvatic transmission. Analysis of YFV cases combined with genomes generated locally using a new protocol reveals an early phase of sylvatic YFV transmission restricted to Minas Gerais, followed in late 2016 by a rise in viral spillover to humans, and the southwards spatial expansion of the epidemic towards previously YFV-free areas. Our results establish a framework for monitoring YFV transmission in real-time, contributing to the global strategy of eliminating future yellow fever epidemics.

---

Yellow fever (YF) is an acute viral hemorrhagic disease responsible for 29000–60000 deaths annually in South America and Africa (*1*) and is the most severe mosquito-borne infection in the tropics (*2*). Despite the existence of a highly effective YF vaccine since 1937, an estimated >400 million unvaccinated people live in areas at risk of infection (*3*). Yellow fever virus (YFV) is a member of the *Flaviviridae* family and is classified into four genotypes: East African, West African, South American I, and South American II (*4*-*8*). YFV transmission occurs mainly via the so-called “sylvatic cycle”, in which non-human primates (NHP) are infected by the bite of infected tree-dwelling mosquitoes, such as *Haemagogus* spp. and *Sabethes* spp. (*9*, *10*). YFV transmission can also occur via a “domestic cycle”, in which humans are infected by *Aedes* sp. mosquitoes that mostly feed on humans (*11, 12*).

Beginning in 2016, the Americas have experienced the highest number of YF human cases and epizootics for decades. Brazil accounts for nearly a quarter of YF cases in the Americas and has reported 1299 confirmed cases and 421 deaths from YF infection since Jul 2016. Notably, the last case of YF in Brazil attributed to a domestic cycle was in 1942. An intensive eradication campaign eliminated *Aedes aegypti* and YF from Brazil in the 1950s (*13*) but the vector became re-established in the 1970s and *Aedes spp.* mosquitoes are now abundant across most of Brazil (*14*). The consequences of a re-ignition of domestic cycle transmission in Brazil would be very serious, as an estimated 35 million people living in areas at risk for YFV outbreaks in Brazil remain unvaccinated against the disease (*3*). New surveillance and analytical approaches are therefore urgently needed to monitor this risk in real-time.

## Yellow fever virus outbreak in Brazil, 2016–2017

The first human case of the current outbreak in Minas Gerais was confirmed in Dec 2016. Between then and the end of Jul 2017, there were 777 PCR-confirmed human cases across 10 states in Brazil, mostly in Minas Gerais (60% of cases), followed by Espírito Santo (32%), Rio de Janeiro (3%), and São Paulo (3%) (*15*). The fatality ratio of severe cases during this epidemic was estimated at 34.5%, comparable to previous outbreaks (*16, 17*). Despite the magnitude and rapid expansion of the outbreak, little is known about its genomic epidemiology. Further, it is uncertain how the virus is spreading through space, and between humans and NHPs, and analytical insights into the contribution of the domestic cycle to ongoing transmission are lacking.

To characterise the 2017 YFV outbreak in Brazil, we first compare the time series of confirmed cases in humans (n=683) and NHP (n=314) reported by public health institutes in Minas Gerais (MG), the epicentre of the outbreak (**Fig. 1A and B, fig. S1**). The time series are strongly associated (cross-correlation coefficient=0.97; p<0.001). Both peak in late January 2017 and human cases are estimated to lag those in NHP by only 4 days (**table S1**). NHP cases are geographically more dispersed in MG than human cases, which are more concentrated in Teófilo Otoni and Manhuaçu municipalities (**Fig. 1D and E**). Despite this, the number of human and NHP cases per municipality are positively correlated (**Fig. 1F**).

**Fig. 1.**
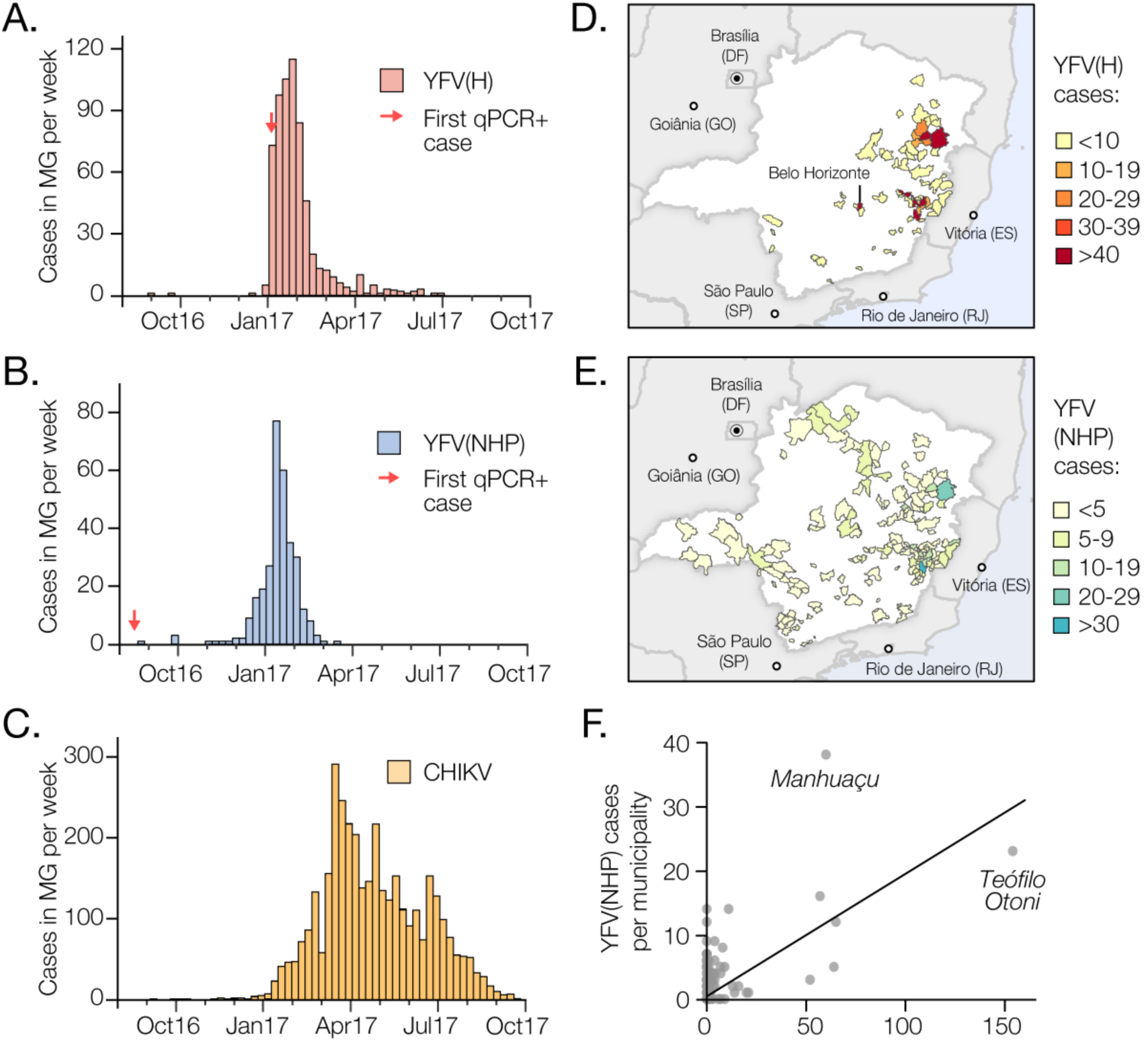
Spatial and temporal epidemiology of YFV in Minas Gerais, 2016-2017. **(A)** Time series of human YFV cases in Minas Gerais confirmed by serology, RT-qPCR, or virus isolation for the first YFV epidemic wave (Aug 2016 to Oct 2017). (**B**) Same as panel A, but for NHP YFV cases, confirmed by RT-qPCR only. (**C**) Same as panel A, but for human CHIKV cases. (**D**) Geographic distribution in Minas Gerais of human YFV cases. (**E**) Geographic distribution in Minas Gerais of NHP YFV cases. **Fig. S2** shows the geographic distribution of CHIKV cases for the same time period. **(F)** Positive association between the number of human and NHP cases in each municipality of Minas Gerais (Pearson’s *r*=0.62; p<0.0001; non-parametric Spearman’s rank *ρ*=0.32; p<0.05). Teófilo Otoni and Manhuaçu are the municipalities with the highest number of human and NHP confirmed YFV cases, respectively.

To establish whether human cases are acquired in proximity to potential sources of sylvatic infection, we estimate the distance between the municipality of residence of each human case and its nearest habitat of potential transmission, determined by using the enhanced vegetation index (EVI) (*18*) (**Materials and Methods**). The average minimum distance between the residence of confirmed human YFV cases and areas with EVI>0.4 is only 5.3km. In contrast, we estimate that a randomly chosen resident of MG lives, on average, at least 51km away from areas with EVI>0.4. Similarly, confirmed human YFV cases reside on average only 1.7km from the nearest NHP case, whereas the mean minimum distance of a randomly chosen resident of MG to the nearest NHP case is 39.1km. This is consistent with the YF infection risk being greatest for people who reside or work in forested areas where sylvatic transmission occurs. Importantly, we find that most human cases (98.5%) were notified in municipalities with YFV vaccination coverage above the threshold of 80% recommended by WHO (*3*) that is sufficient to prevent and control outbreaks. On average, human YFV cases would need to travel 65km from their place of residence to reach an area where vaccination coverage is <80%.

## Risk of YFV domestic transmission

YFV was recently detected in *Ae. albopictus* mosquitoes caught in MG in Jan 2017 (*19*). Further, experiments suggest that *Aedes* spp. mosquitoes from the neighbouring state of Goiânia can transmit Brazilian YFV, albeit less effectively than vectors from elsewhere in Brazil (*20*). It is therefore important to investigate whether YFV cases in MG occur where and when *Aedes* spp. vectors are active. To do so, we analysed confirmed chikungunya virus (CHIKV) cases from MG in 2016-17 (**Fig. 1C**).

CHIKV is transmitted by the domestic mosquitoes *Ae. aegypti* and *Ae. albopictus* (*21*). There were 3755 confirmed CHIKV cases reported in MG during Jan 2015 to Oct 2017. The CHIKV epidemic in MG in 2017 began later and lasted longer than the YFV outbreak (**Fig. 1C**), consistent with the hypothesis that YF and CHIKV in the region are transmitted by different vector species. However, 26 of the municipalities with YFV human cases also reported CHIKV cases (**Fig. 1D** and **fig. S2**), indicating that YFV *is* present in municipalities with *Aedes* mosquitoes. The mean YFV vaccination rate in districts with both YFV and CHIKV cases is 72.3% (range=61-78%). Thus, relatively high vaccination rates in the locations in MG where YF spillover to humans occurs, and potentially lower vector competence (*21*), may be ameliorating the risk of establishment of a domestic YFV cycle in the state. However, adjacent urban regions (including São Paulo and Rio de Janeiro) have lower YFV vaccination rates (*3*), receive tens of millions of visitor per year (*22*), and have recently experienced YF human cases (*17*). Thus, the possibility of sustained domestic transmission of YF in southern Brazil necessitates continual virological and epidemiological monitoring.

## Epidemiological model to investigate YFV transmission mode

We next sought to establish a framework to evaluate the routes of YFV transmission during an outbreak from the characteristics of infected individuals. Specifically, we assess whether an outbreak is driven by sylvatic *vs.* domestic transmission by comparing the age and sex distributions of observed YFV cases with those expected under a domestic cycle in Minas Gerais. For example, an individual’s risk of acquiring YFV via the sylvatic cycle depends on their likelihood of travel to forested areas, which is typically highest among male adults (*23*). In contrast, under a domestic transmission cycle we expect more uniform exposure across age- and sex-classes.

The male-to-female sex ratio of reported YFV cases in MG is 5.7 (*i.e*. 85% of cases are male) and incidence is highest among males aged 40-49 (**Fig. 2**). We compare this distribution to that expected under two models of domestic cycle transmission. In model M1, all age- and sex-classes vary in their vaccination status but are equally exposed to YFV, a scenario that is typical of arboviral transmission (*24*). Under model M1, predicted cases are characterized by a sex ratio ∼1 and incidence peaks among individuals aged 20-25 (**Fig. 2**). In model M2, we assume that the pattern of YFV exposure among age- and sex-classes follows that observed for CHIKV. The sex ratio of reported CHIKV cases in MG is 0.49 (*i.e.* 33% of cases are male; **fig. S3**). Under model M2, predicted incidence is highest in females aged >30. The discrepancy between the observed age distribution and that predicted under the two domestic cycle models indicates that the YF epidemic in MG is dominated by sylvatic transmission. The method introduced here shows that age- and sex-structured epidemiological data can be used to qualitatively evaluate the mode of YFV transmission during an outbreak.

**Fig. 2.**
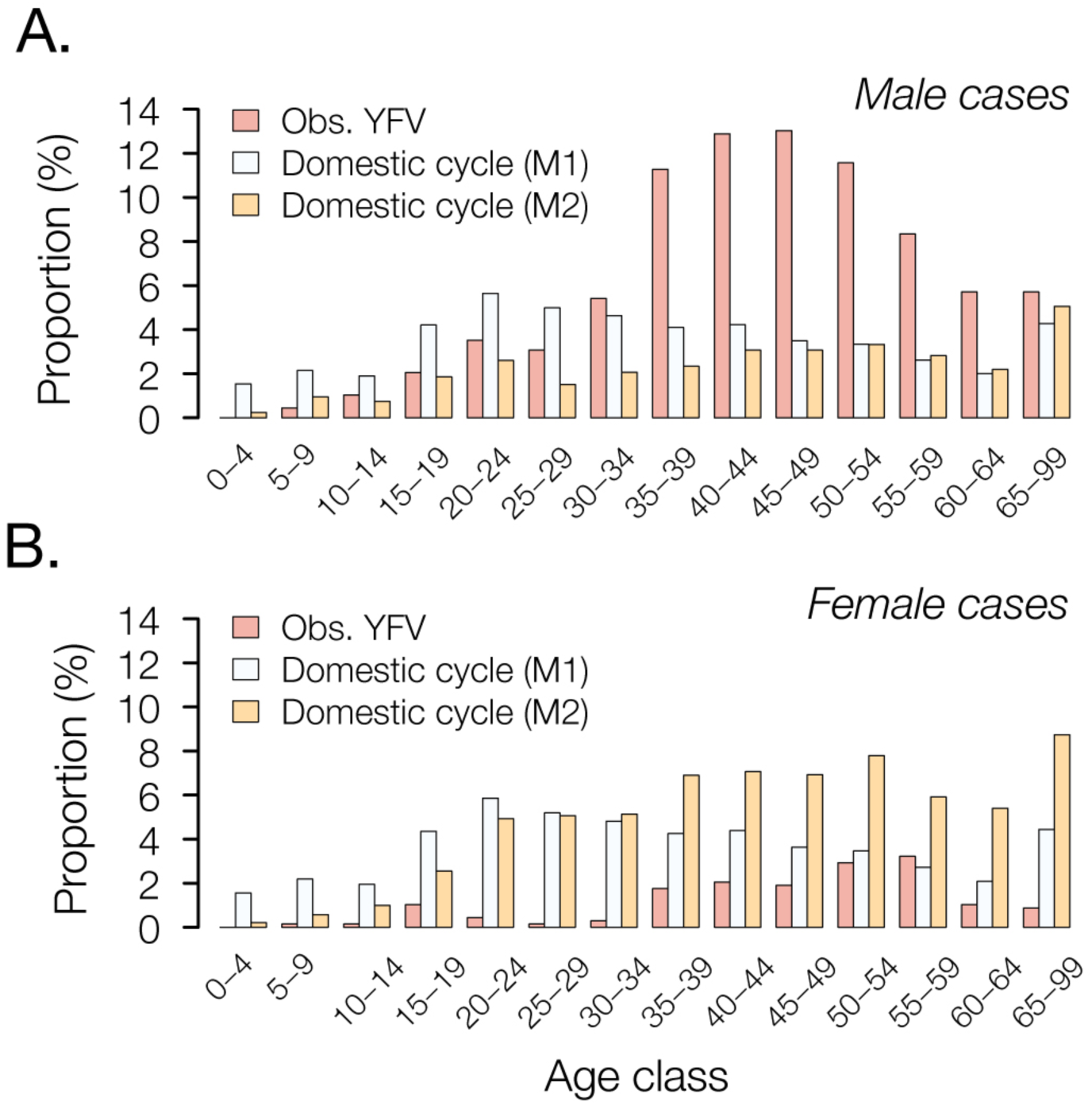
Age and sex distribution of YFV cases in Minas Gerais, 2016-2017. The red bars show the proportion of observed YFV cases in Minas Gerais that occur in each age class, in males (panel **A**) and females (panel **B**). These empirical distributions are notably different from those predicted under two models of domestic cycle transmission. Model M1 (white bars) assumes that all age- and sex-classes are equally exposed to YFV but vary in vaccination status, whilst model M2 (orange bars) assumes that variation in YFV exposure among classes follows that observed for CHIKV (**fig. S3**; **Materials and Methods**).

## Genomic surveillance of the Brazilian YFV outbreak

During a YF outbreak it is important to undertake virological surveillance to (*i*) track epidemic origins and transmission hotspots, (*ii*) characterise genetic diversity to aid molecular diagnostics, (*iii*) detect viral mutations associated with disease severity, and (*iv*) exclude the possibility that human cases are caused by vaccine reversion.

We generated and analyzed 52 complete YF genomes from infected humans (*n*=32) and non-human primates (*n*=20) from the most affected Brazilian states, including Minas Gerais (*n*=40), Espírito Santo (*n*=8), Rio de Janeiro (*n*=2), São Paulo (*n*=1) and Bahia (*n*=1) (**Fig. 3, tables S2, S3**). We included two new genomes from samples collected in 2003 during a previous YFV outbreak in MG in 2002–2003 (*25*). Although genomes were generated using a combination of sequencing methods (**table S2**), 70% of the new sequences were generated in Minas Gerais itself using a rapid MinION portable sequencing protocol for YFV (*26*) (**table S4**). This protocol was made publicly available in May 2017 following pilot sequencing experiments using a cultured vaccine strain (see **Materials and Methods**). Median genome coverages were similar for samples obtained from NHP (99%; median Ct = 11) and from human cases (99%; median Ct=15) (**table S5**).

**Fig. 3.**
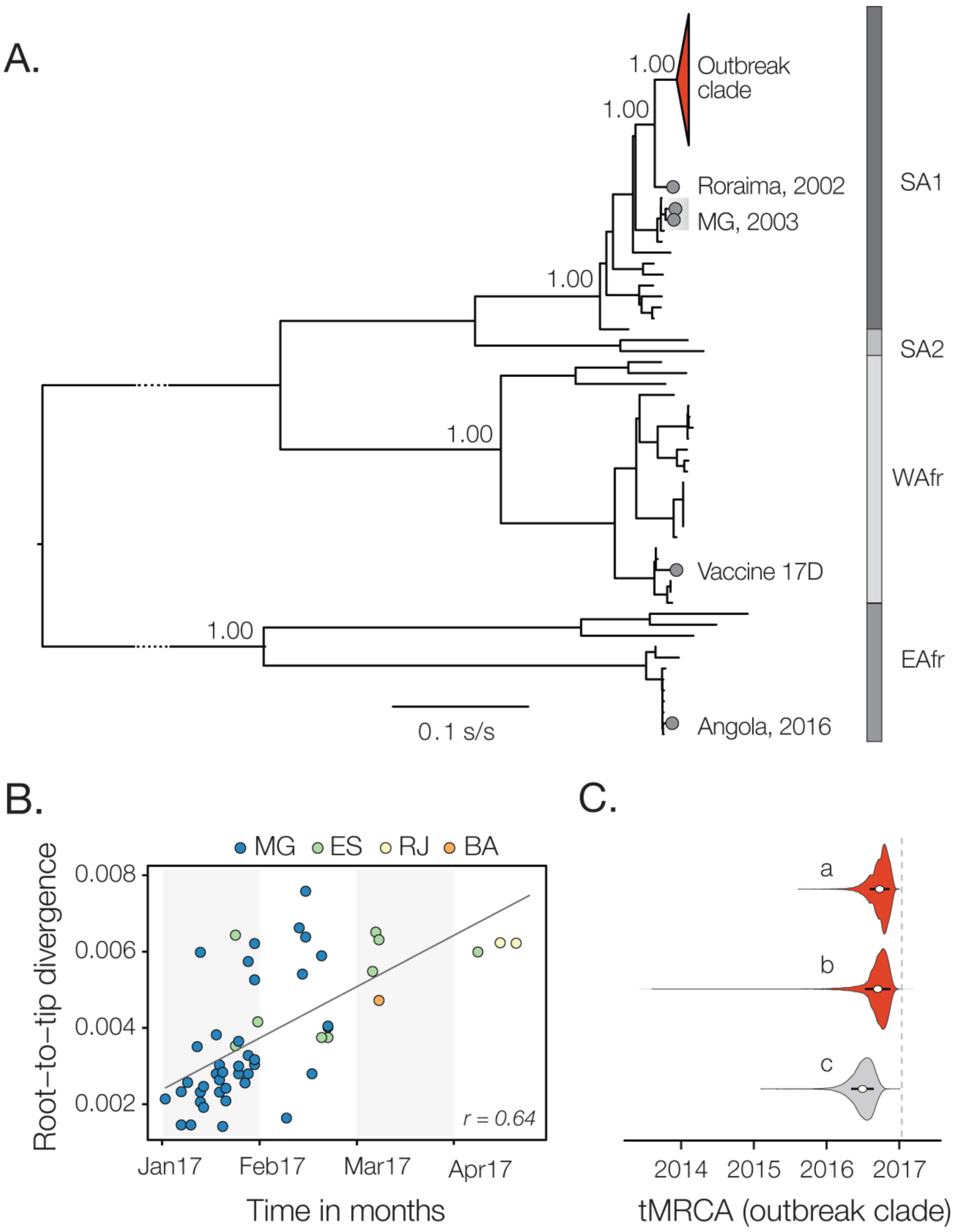
Genomic surveillance of the Brazilian YFV epidemic. **(A)** Maximum likelihood phylogeny of complete YFV genomes (*n*=103) shows that all sequences from the outbreak belong to a single strongly supported clade within the South America I (SA1) genotype (red triangle; see **fig. S5** for an annotated tree and **Fig. 4** for details of the outbreak clade). The scale bar is in units of substitutions per site (s/s). Highlighted sequences (grey circles) represent (i) the most closely related genome to the outbreak, sampled from a human isolated collected in 2002 in Alto Alegre, Roraima, North Brazil, (ii) two historical Minas Gerais sequences collected in 2003, (iii) the vaccine strain 17D used in Brazil, and (iv) the YFV outbreak Angola in 2015-2016 (*12*). (**B**) Root-to-tip regression of sequence sampling time against genetic divergence from the root of the outbreak clade. Sequences are coloured according to sampling location. (**C**) Violin plots showing estimated posterior distributions (white circle=mean) of the time of the most common ancestor (tMRCA) of the outbreak clade. Estimates were obtained using two different datasets (grey=SA1 genotype, red=outbreak clade only) and under different evolutionary models: a=uncorrelated lognormal relaxed clock (UCLN) model with a skygrid tree prior with covariates; b=UCLN model with a skygrid tree prior without covariates; c=fixed local clock model (see **Materials and Methods**).

To put the newly generated YFV genomes in a global context, we added our genomes to 61 publicly available genomes, including two NHP genomes available from the ongoing outbreak (*27*). We developed and applied a new automated online phylogenetic tool to reliably identify and classify YFV gene sequences, which is also publicly available (see **Materials and Methods**; **fig. S4**). Phylogenies estimated using maximum likelihood and Bayesian methods, and the genotyping tool, all consistently place the Brazilian outbreak strains in a single clade within the South America I (SA1) genotype (*27*), with maximum statistical support (bootstrap=100%; posterior probability>0.99) (**Fig. 3A**; **fig. S4**).

The outgroup to the YFV outbreak clade is strain BeH655417, sampled in Roraima, north Brazil in 2002. In contrast, local isolates sampled during the previous outbreak in MG in 2003 are more distantly related to the outbreak clade within the SA1 genotype (**Fig. 3**). Thus the 2017 outbreak was more likely caused by a YFV strain introduced from northern Brazil (or another unsampled region) than by the re-emergence of a lineage that had persisted in MG, although low sampling densities mean this conclusion is provisional. The 14-year gap between the current outbreak and the date of the most closely related non-outbreak strain agrees with the reported periodicity of YF outbreaks in northern Brazil (*28*), which is thought to be dictated by the rate of accumulation of susceptible NHP hosts (*16, 29*).

At least 7 PCR-confirmed YFV human cases in MG received a YF vaccine up to 3 days before onset of symptoms. To test the scenario that these infections were caused by natural infection, and not by vaccine reactivation, we sequenced the YFV genomes of two of these cases (**Fig. 3A, table S3**). Our phylogenetic analysis (**Fig. 3A**) clearly shows that the human cases represent natural infections caused by the ongoing outbreak, and are conclusively not derived from the 17D vaccine strain which belongs to the West African YFV genotype (**Fig. 3A**).

## Unifying YFV epidemiology and molecular evolution

Viral genomes are a valuable source of information about epidemic dynamics (e.g. (*30*)) but have rarely been used to investigate YFV outbreaks in detail. Here we show how a suite of three analytical approaches, that combine genetic, epidemiological and spatial data, can provide high-resolution insights into YFV transmission.

First, we used a Bayesian method (*31*) to explore potential covariates of fluctuations in the effective population size of the YFV outbreak in 2017. After confirming that genetic divergence in the YFV outbreak clade accumulates over the timescale of sampling (**Fig. 3B, fig. S5**), we tested which epidemiological time series best describe trends in inferred YFV effective population size. Our analysis reveals that effective population size fluctuations of the YFV outbreak are almost equally well explained by the dynamics of human and NHP YFV cases (inclusion probability=0.55 for human cases and =0.45 for NHP cases). These two YFV time series explain the genetic diversity dynamics of the ongoing outbreak 10^3^ times better than the CHIKV time series, whose inclusion probability is <0.001 and which represents viral transmission by *Aedes* spp. vectors. One benefit of this approach is that epidemiological data contribute to estimation of the timescale of the outbreak. By incorporating the time series of YFV cases into evolutionary inference, we estimate the time of the most recent common ancestor (TMRCA) of the outbreak clade to be late-Sep 2016 (95% Bayesian Credible Interval, BCI: Jun-Dec 2016) (**Fig. 3C, fig. S6**), consistent with the date of the first PCR-confirmed case of YFV in NHP in MG (Jul 2016). The uncertainty around the TMRCA estimate is reduced by 22% when epidemiological and genomic data are combined, compared to genetic data alone (**Fig. 3C**).

Second, in order to better understand YFV transmission between humans and NHP we reconstructed and measured the movement of YFV lineages between the NHP reservoir and humans, using a phylogenetic structured coalescent model (*32*). Although previous studies have confirmed that YFV is circulating in five neotropical NHP families (Aotidae, Atelidae, Callitrichidae, Pitheciidae, Cebidae; **Fig 4E;**, **table S3**), thus far NHP YFV genomes during the 2017 outbreak have been recovered only from *Alouatta* spp. (family Cebidae) (*27*). In this analysis we used the TMRCA estimate obtained above (**Fig. 3C**) to inform the phylogenetic timescale (**Fig 4B**). Even though most (61%) YFV genomes were from human cases, almost all internal nodes in the outbreak phylogeny whose host state is well supported (posterior probability >0.8) are inferred to belong to the NHP population, consistent with an absence of domestic transmission and in agreement with the large number of NHP cases reported in southeast Brazil (*17*). Only one internal node (the ancestor of sequences from human cases M26 and M98) is assigned as human with a posterior probability >0.8 (**Fig. 4A**). Travel information for our YFV human cases (**table S3**) shows that M26 was living in a rural area of São Caetano do Sul, SP, located >1000km from the urban area of Teófilo Otóni, MG, where M98 was resident. Thus, these two human cases are unlikely to be epidemiologically linked; their ancestor has been likely incorrectly assigned due to the relative under-representation of YFV sequences from NHP compared to human sequences in our data. Despite this, the structured coalescent approach clearly reveals significant changes in the frequency of NHP-to-human host transitions through time, rising from zero around Nov 2016 and peaking in Jan 2017 (**Fig. 4B**). Remarkably, this phylogenetic trend matches the time series of confirmed YFV cases in MG (**Fig. 1B**), demonstrating that viral genomes, when analysed using appropriate models, can be used to quantitatively track the dynamics of zoonosis during the course of a complex outbreak (*33*).

**Fig. 4.**
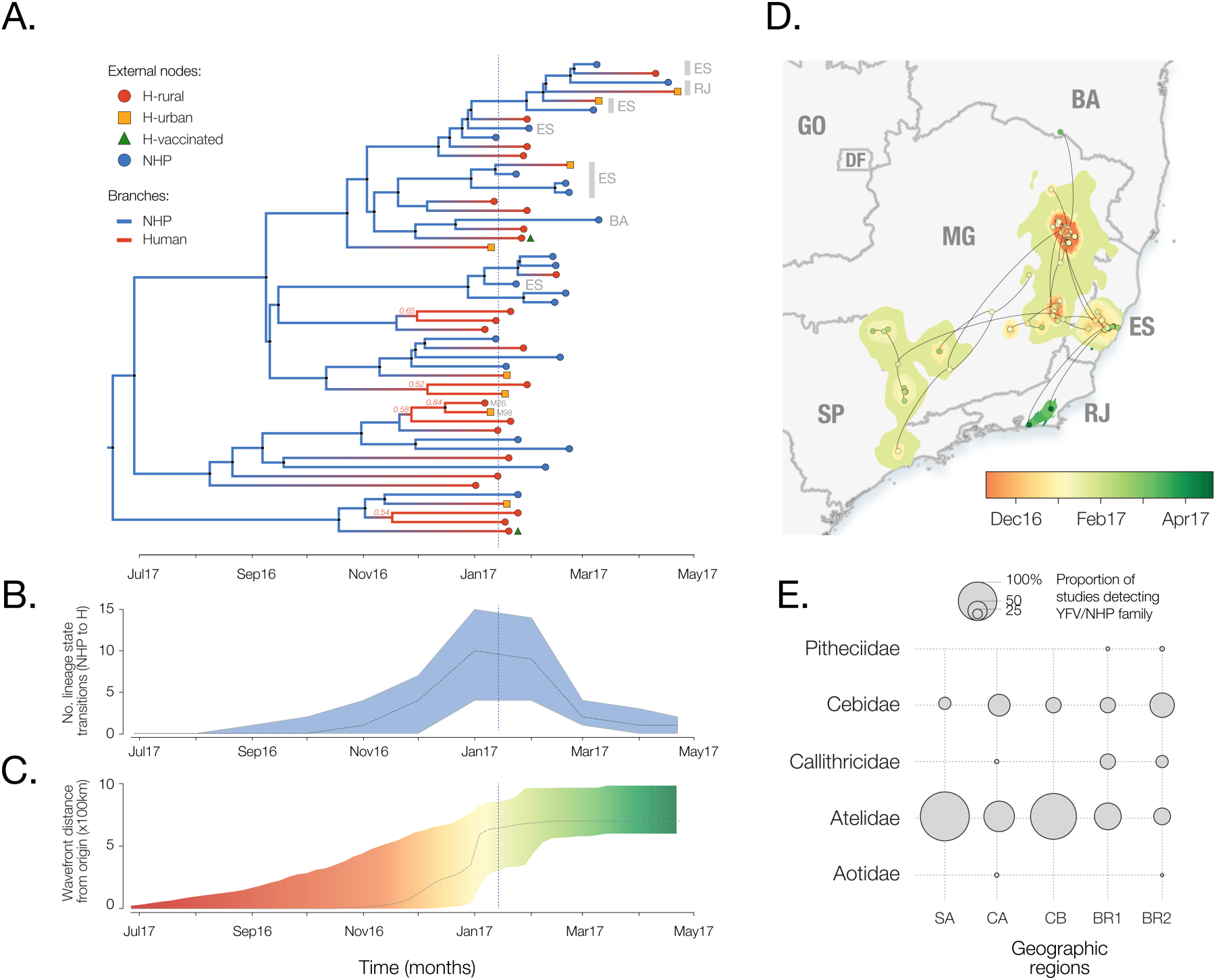
Spatial and evolutionary dynamics of YF in southeast Brazil. **(A)** Maximum clade credibility phylogeny inferred under a two-state (human and NHP) structured coalescent model. External node symbols denote sample type. Grey bars and labels to the right indicate sample location (RJ=Rio de Janeiro, ES=Espírito Santo, BA=Bahia; other sequences were sampled in MG). Internal nodes whose posterior state probabilities (human or NHP) are >0.8 are annotated by circles. Internal branches are coloured blue for NHP, red for human. A fully annotated tree is shown in **fig. S7 (B)**. The average number of YFV phylogenetic state transitions (from NHP to human) per month (see also panel A). Solid line=median estimate, shaded area=95% Bayesian Credible Interval (BCI). (**C**) Expansion of the YFV epidemic wavefront estimated using a continuous phylogeographic approach (*35*). For each point in time, the plot shows the maximum spatial distance between phylogeny branches and the inferred location of outbreak origin. Solid line = median estimate, shaded area = 95% BCI. **(D)** Reconstructed spatiotemporal diffusion of the YFV outbreak. Phylogeny branches are arranged in space according the locations of phylogeny nodes (circles). Locations of external nodes are known, whilst those of internal nodes are inferred (*34*). GO=Goiás, SP=São Paulo. Shaded regions show 95% credible regions of internal nodes. Nodes and uncertainty regions are coloured according to time. (**E**) Summary of the frequency of detection of YFV in non-human primates in the Americas (*35*). Circle sizes represent the proportion of published studies (*n*=15) that have detected YFV in each primate family and region (**table S3**). SA=South America region except Brazil, CA=Central America, CB=Caribbean, BR1=Brazil (before 2017), BR2=Brazil (this study).

Third, we measured the outbreak’s spatial spread using a phylogenetic relaxed random walk approach (*36*) (see **Materials and Methods**; **table S6**). When projected through space and time (**Fig. 4D**; **Movie S1**), the outbreak phylogeny reveals a southerly dissemination of virus lineages from their inferred origin in northeast MG, towards densely populated areas, including Rio de Janeiro and São Paulo (where YF vaccination was not recommended until Jul 2017 and Jan 2018, respectively). We estimate virus lineages move on average 7.5 km/day (95% BCI: 3.8 to 21.8 km/day) (*34*). This velocity remains high when human YFV terminal branches are removed (8.2 km/day) and therefore reflects most likely the rate of YFV lineage movement within the sylvatic cycle and not the movement of asymptomatic infected humans. These rates are higher than expected given the distances typically travelled by NHPs in the region (*37*), and suggest the possibility that movement of some YFV lineages may have been aided by human activity, e.g. by transport of infected mosquitoes in vehicles (*38*) or hunting or illegal trade of NHPs in the Atlantic forest (*39*, *40*). The epidemic wavefront (maximum distance of phylogeny branches from the inferred epidemic origin) moved little before Nov 2016, after which the wavefront expanded at ∼7km/day, until Feb 2017. Thus by the time YF was declared a public health emergency in MG (13 Jan 2017; dashed lines in **Figs. 3c and 4a-c**), the epidemic had already travelled >500km from northeast MG (**Figs 4c,d**) and caused >100 cases in both humans and NHP (**Fig. 1**). Notably, the first detection in humans in Dec 2016 was concomitant with both the spatial expansion of the epidemic (**Fig 4c**) and the rise in the number of NHP-to-human zoonoses (**Fig. 4B**), most probably driven by an increase in the abundance of sylvatic vectors. Thus, the outbreak lineage appeared to circulate among NHP in a geographically restricted area for several months before human cases were detected.

## Conclusion

Epidemiological and genomic surveillance of human and animal populations at risk is crucial for the early detection of YFV transmission and its rapid containment. The upsurge of cases since Dec 2017 means the YFV epidemic in Brazil is continuing to unfold. Longitudinal studies of NHP are needed to understand how YF lineages disseminate across South America between YF outbreak years, and how YF epizootics might be determined by the dynamics of susceptible animals in the reservoir. To achieve the World Health Organization’s goal to eliminate yellow fever epidemics by 2026, YF surveillance demands a global, coordinated strategy. Our results and analyses show that rapid genomic surveillance of YFV in NHP and humans, when integrated with epidemiological and spatial data, can help anticipate the risk of human YFV exposure through space and time and monitor the likelihood of sylvatic versus domestic transmission. We hope that the toolkit introduced here will prove useful in guiding the control of yellow fever outbreaks in a resource-efficient manner.

## Acknowledgements

This work supported in part by CNPq # 400354/2016-0 and FAPESP # 2016/01735-2. N.R.F. is supported by a Royal Society and Wellcome Trust Sir Henry Dale Fellowship (204311/Z/16/Z), internal HEFCE GCRF grant 005073, and John Fell Research Fund Grant 005166. This research received funding from the ERC under grant agreement 614725-PATHPHYLODYN and from the Oxford Martin School. MUGK is supported by the Society in Science, the Branco Weiss Fellowship, administered by ETH Zurich and acknowledges funding from a Training Grant from the National Institute of Child Health and Human Development (T32HD040128) and the National Library of Medicine of the National Institutes of Health (R01LM010812, R01LM011965). SD is funded by the Fonds Wetenschappelijk Onderzoek (FWO, Flanders, Belgium). GB acknowledges support from the Interne Fondsen KU Leuven / Internal Funds KU Leuven. ACdC is funded by FAPESP # 2017/00021-9. ACdC and ECS are grateful to Illumina, Zymo Research, Sage Science and Promega for donation of reagents. BBN and SC are supported by the EU’s Horizon 2020 Programme through ZIKAlliance (grant 734548), the Investissement d’Avenir program, the Laboratoire d’Excellence Integrative Biology of Emerging Infectious Diseases program (grant ANR-10-LABX-62-IBEID), the Models of Infectious Disease Agent Study of the National Institute of General Medical Sciences, the AXA Research Fund, and the Association Robert Debré. PL and MAS acknowledge funding from the European Research Council (grant agreement 725422-ReservoirDOCS) and from the Wellcome Trust Collaborative Award 206298/Z/17/Z. PL acknowledges support by the Research Foundation, Flanders (Fonds voor Wetenschappelijk Onderzoek, Vlaanderen, G066215N, G0D5117N and G0B9317N).

## Materials and Methods

### Description of epidemiological data

Between Jan 2015 and Sep 2017, 2571 samples from patients residing in 212 municipalities of Minas Gerais (MG) with symptoms compatible with YFV infection were tested at the Fundação Ezequiel Dias (FUNED), located in Belo Horizonte, MG, southeast Brazil (**Fig. S1**). During the same period, 9555 human samples from patients residing in 362 municipalities of MG were tested for CHIKV infection in the same laboratory (**Fig. S1**). Following Pan American World Health Organization (PAHO) guidelines, YFV human samples were obtained ≤ 6 days after the onset of clinical symptoms, after which they were subjected to RT-qPCR. If samples were obtained >6 days after onset of disease, serological confirmation of YFV through IgM detection was performed. Due to potential cross-reactivity of serological assays, a positive YF serological test performed 6 days after onset of symptoms can indicate either recent YF infection, past YF vaccination, or infection with other circulating flaviviruses, such as Zika virus or dengue virus (*41*). Sex, age, municipality of residence, date of sample collection, and date of onset of symptoms were available for human YFV cases, YFV(H). The post-mortem liver tissue samples from non-human primates (NHP) in MG reported here were tested using YFV RT-qPCR at the Universidade Federal Minas Gerais (UFMG) in Belo Horizonte. The information available for the YFV-positive NHP cases included primate family or species, date of capture, and municipality of sample collection.

**Fig. S1.**
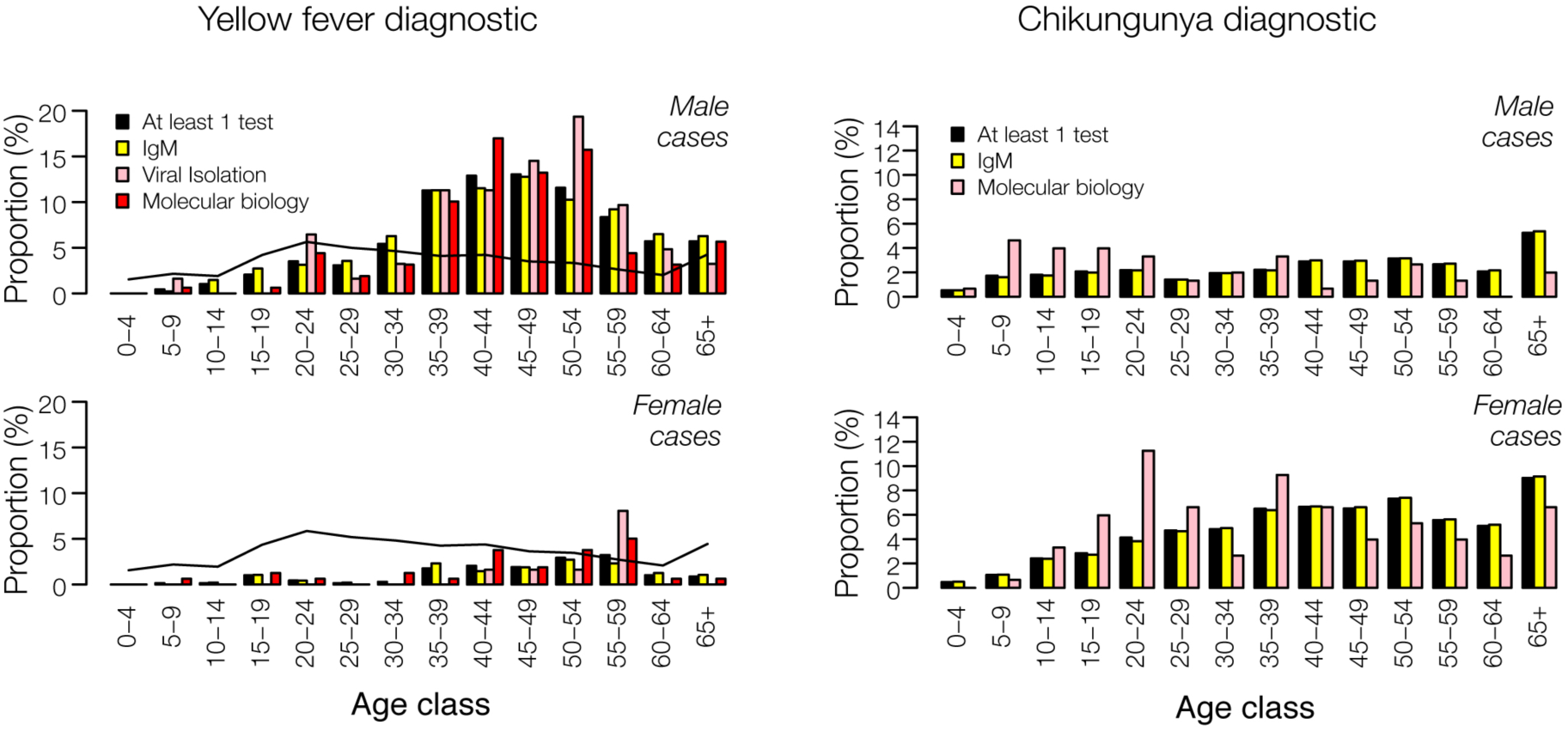
Sensitivity of diagnostics for human YFV (left) and CHIKV (right) samples in Minas Gerais. The proportion of samples positive for at least one test (black bars) in each age- and sex-class is shown, together with the proportion of positives for each test separately (IgM, viral isolation, or RT-qPCR). The black line indicates the average proportion of cases per age class, for male and female cases.

From the laboratory cases tested in MG between Jan 2015 and October 2017, the following datasets were prepared for use in epidemiological analyses. Note that the same patient may have been tested with different tests, so the sample size (N) given below equals the total number of individuals with at least one positive test, which may be less than the sum of the number of positive tests:

- <u>Dataset A:</u> YFV(H) cases confirmed either by RT-qPCR (n=159) or by virus isolation (n = 62) or by IgM (n=478) at FUNED (n=683);
- <u>Dataset B:</u> YFV(NHP) cases confirmed by RT-qPCR in liver tissue analysed at the UFMG (N=314);
- <u>Dataset C:</u> CHIKV confirmed by RT-qPCR (n=144) or by IgM (n=3609) at FUNED (N=3755; no virus isolation was performed for CHIKV).
- <u>Dataset D:</u> YFV(H) confirmed either by RT-qPCR (n=159) or by virus isolation (n=62) at FUNED (n=221);
- <u>Dataset E:</u> CHIKV confirmed by RT-qPCR at FUNED (n=144).

The geographic distribution of YFV(H), YFV(NHP) and CHIKV cases are shown in **Fig. 1D**, **Fig. 1E** and **fig. S2**, respectively. Note that these maps correspond, respectively, to datasets A, B and C described above.

**Fig. S2.**
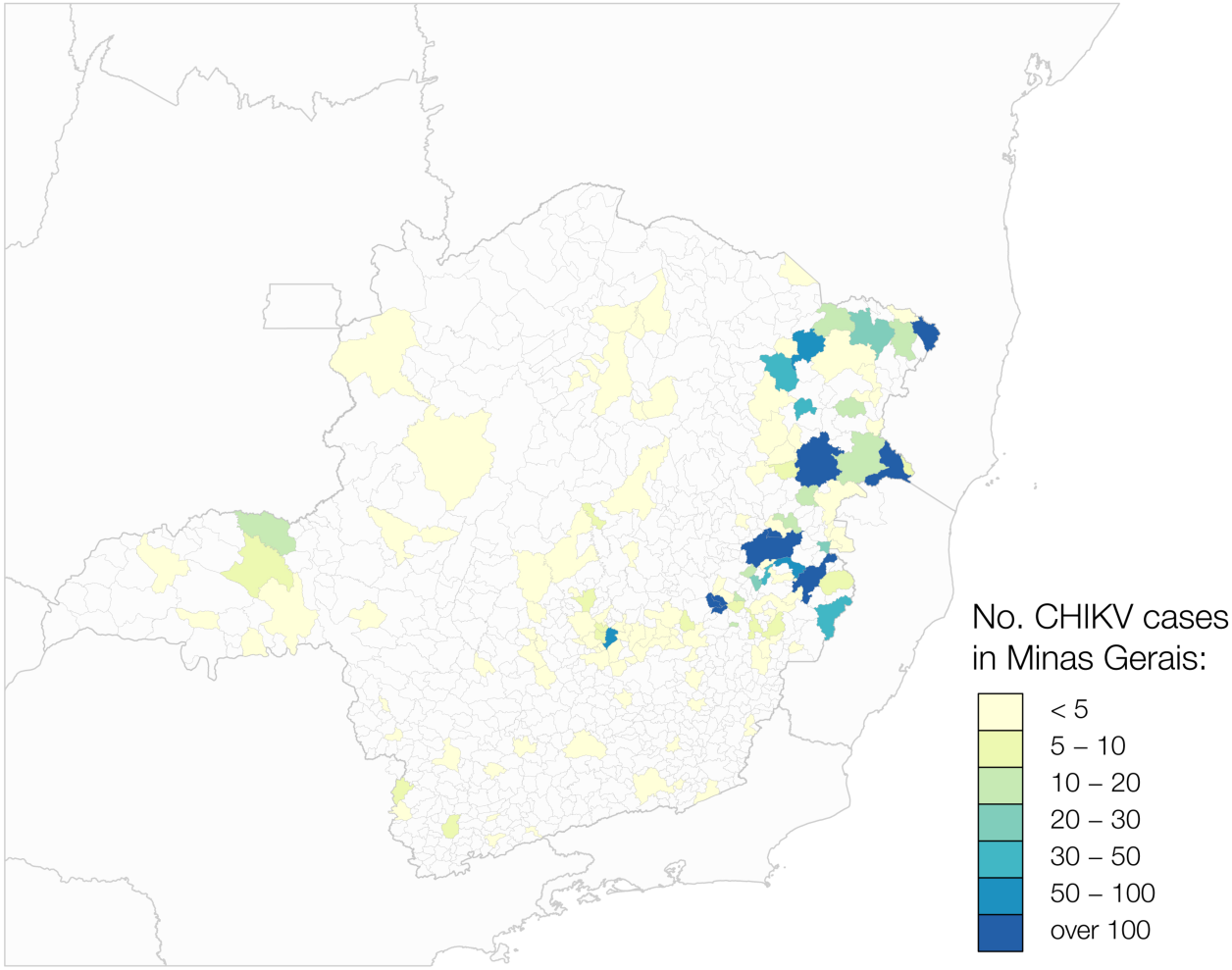
Geographic distribution of CHIK cases in Minas Gerais. The figure shows cases confirmed by serology, RT-qPCR, or virus isolation in Minas Gerais from Jan 2015 to October 2017 (corresponding to dataset C).

To assess the association between the time series of YFV(NHP) and YFV(H) cases, we computed pairwise cross correlations among datasets A, B, and C, correcting for time lag and assuming that each dataset followed a unimodal distribution across time that covered a single epidemic wave of YFV. The correlations and corresponding P-values are shown in **table S1**.

**Table S1.** Time-series cross correlation analysis. A, B and C correspond to datasets A, B and C described in the text. Period: time frame during which the comparison is undertaken. The dataset in brackets is fixed during that period, while the other dataset is shifted temporally to correct for potential lag. *p*-value: *p*-value of the correlation between the two “auto-correlated” series, accounting for the time-lag. Time lag: the lag between the two series estimated to nearest day via linear interpolation. Similar observations were obtained when datasets D and B, D and C, and B and E were compared (data not shown).

| Comparison | A vs B | A vs C | B vs C |
| --- | --- | --- | --- |
| Period | 1 Aug 2016 to<br>1 Oct 2017 (A) | 1 Aug 2016 to<br>1 Oct 2017 (A) | 1 Aug 2016 to<br>1 Oct 2017 (A) |
| *P*-value | <0.0001 | <0.0001 | <0.0001 |
| Cross-correlation | 0.971 | 0.757 | 0.725 |
| Time lag | B is 4 days<br>ahead of A | C is 62 days<br>behind A | C is 64 days<br>behind B |

### Model of age-sex distributions under domestic and sylvatic transmission cycles

To characterize whether human YF cases result from a domestic or sylvatic transmission cycle we examined the age-sex distribution of human YF cases in MG between Dec 2016 (the date of first confirmed human YFV RT-qPCR case) and October 2017 (see **Fig. 1**). Two models were developed to predict the age-sex distribution of YFV cases expected under a domestic cycle. In *model M1*, we assume that exposure to YFV in the domestic cycle is independent of sex and age. We reconstructed the resulting age-sex distribution from the underlying population age pyramid in MG (*42*) and from vaccine coverage per birth cohort (*3*). The expected number of individuals of age *a* and sex *s* that are at risk of YFV infection is then:

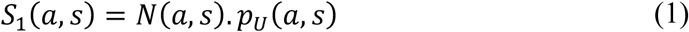

where *N*(*a*, *s*) is the number of individuals of age *a* and sex *s* in the population and *p_U_*(*a*, *s*) is the proportion of unvaccinated individuals in that group. We assume that the proportion of vaccinated individuals is independent of sex in a given birth cohort. The expected proportion of YFV cases that are of age *a* and sex *s* is therefore:

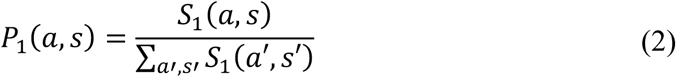

In *model M2*, we assume that, under a scenario of domestic cycle transmission, the risk of exposure to YFV for a susceptible individual would be proportional to that seen for CHIKV cases (**fig. S3**).

**Fig. S3.**
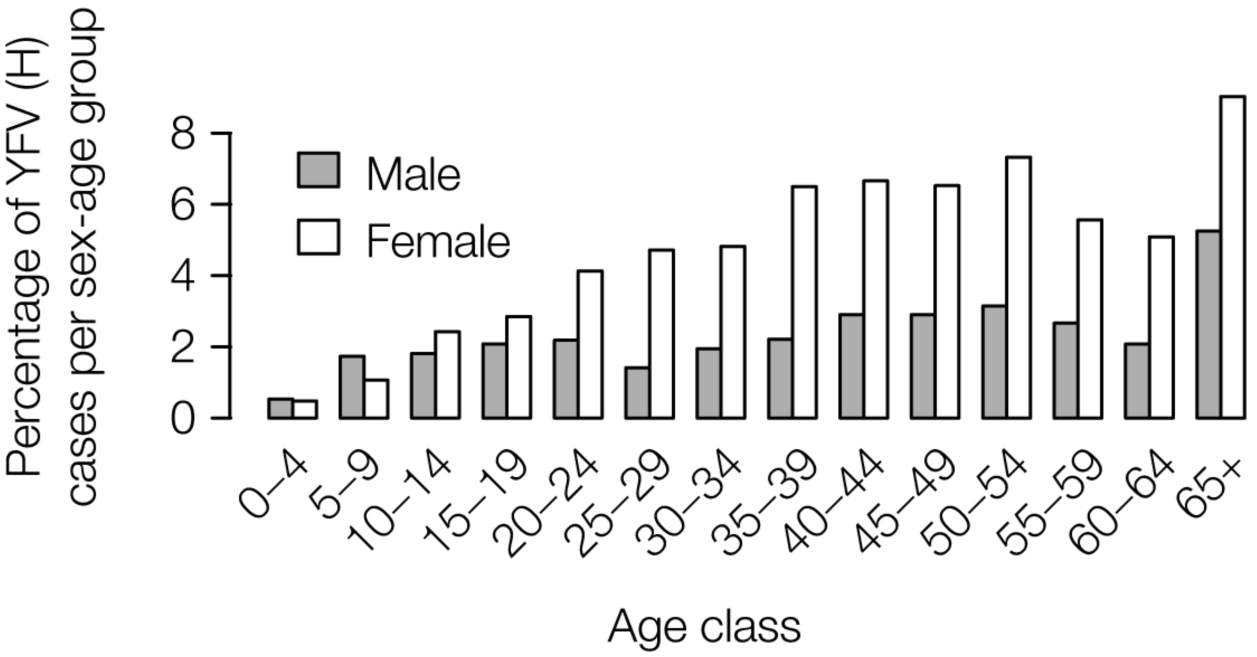
Age-sex distribution of reported CHIKV cases in Minas Gerais. The y-axis shows the percentage of CHIV cases belonging to each age- and sex-class that were confirmed by at least one diagnostic test in Minas Gerais between Jan 2015 and October 2017 (dataset C).

Let *C*(*a*, *s*) denote the number of reported CHIKV cases of age *a* and sex *s*. For an individual of age *a* and sex *s*, the relative risk of being reported as a CHIKV case is defined as:

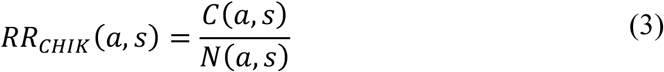

Thus in model M2, the expected proportion of YFV cases that are of age *a* and sex *s* is:

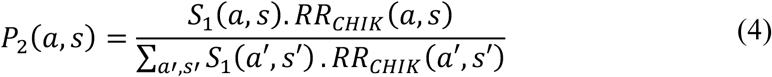

### Estimating expected spatial distances to the source of YF infection

Many human YFV cases were reported in cities across the region and the travel history of most cases remains unknown. To assess the likelihood of a sylvatic transmission cycle scenario, in which most infections occur in forested areas, we calculate the average great circle distance between the place of residence of each human case and the nearest location with environmental conditions suitable for sylvatic transmission (*43*). We then compare this distance to that expected for typical (non-YFV infected) residents of Minas Gerais, estimated using high-resolution population datasets from 2015 (*44*). We used overall greenness of the environment [Enhanced Vegetation Index (EVI) (*45*)] to identify areas with environmental conditions suitable for sylvatic transmission. The EVI has been previously determined to be the best fitting predictor of seasonal YFV transmission (*46*). Several thresholds of EVI for each municipality were considered: 0.33 (5%), 0.41 (50%), 0.46 (95%). Great-circle distances were calculated using the “rdist.earth” function in R (*47*). We also calculated the distance to areas with known occurrences of positive non-human primates, again for both confirmed YF cases in humans and for typical residents of Minas Gerais.

### Ethical statements for biological data

The project was supported by the Pan American World Health Organization (PAHO) and the Brazilian Ministry of Health as part of arboviral genomic surveillance efforts. Human samples were previously obtained for routine diagnostic purposes from persons visiting local clinics in Minas Gerais and Rio de Janeiro. Residual anonymized clinical diagnostic samples, with no or minimal risk to patients, were provided for research and surveillance purposes within the terms of Resolution 510/2016 of CONEP (Comissão Nacional de Ética em Pesquisa, Ministério da Saúde; National Ethical Committee for Research, Ministry of Health). We included 121 samples extracted at the Fundação Ezequiel Dias (FUNED), the main central public health laboratory in Minas Gerais (MG) (*sub*-*study I*). An additional 8 non-human primate (NHP) samples were extracted at the Universidade Federal de Minas Gerais (UFMG) and subsequently sent to FIOCRUZ Bahia for sequencing. Human samples were also processed at the Reference Centre for Arbovirus in Rio de Janeiro, the Laboratory of Flavivirus at FIOCRUZ Rio de Janeiro (*sub*-*study II*). Ethical approval for human samples was obtained from CEP/CAAE: 0026.0.009.000-07, with Institutional Review Board approval numbers 027/2007 and 1.920.256). Samples obtained from the Reference Centre for Arbovirus of São Paulo, Adolfo Lutz Institute (IAL) have been processed in agreement with routine surveillance activities from the Brazilian Ministry of Health and under the CEUA (Comitê de Ética de Uso de Animas em Pesquisa) registration number 02/2011.

**Table S2.** Laboratories involved in YFV genome sequencing. H=Human, NHP=Non-human primate, MG = Minas Gerais, BA=Bahia, RJ=Rio Janeiro, ES=Espírito Santo.

| **Institution, State** | **Sample collection** | **Sequencing Platform** | **Sub-study** |
| --- | --- | --- | --- |
| FUNED, MG | MG, SP | MinION | I |
| FIOCRUZ, BA | MG, BA | MinION | II |
| FIOCRUZ, RJ | MG, ES, RJ | Ion Torrent™ | III |
| IAL, SP | ES | MiSeq | IV |

### Viral RNA isolation and sample processing

Human clinical samples included tissue and serum or plasma. In brief, viral RNA was extracted from 200 μL of clinical sample using QIAmp Viral RNA Minikit (Qiagen) according to the manufacturer’s instructions but with several protocol changes. Tissue samples were first homogenised using a TissueLyser LT (Qiagen). A small piece of tissue (∼2 mm diameter) was cut using a disposable scalpel and added to a 2 mL Eppendorf tube containing a 5 mm stainless steel bead (Qiagen). 560 μl AVL lysis buffer (Qiagen) was added to each tube and the sample was homogenised for 5 min at 50 Hz on a TissueLyser LT followed by a 10 min incubation at room temperature to lyse virions. Samples were centrifuged at 1,200g for 2 min to pellet cellular material, and 500 μL of supernatant was transferred to a new tube containing 500 μL of 100% EtOH. For serum or plasma samples, 200 μL of the sample was added to 560 μL of AVL lysis buffer (Qiagen) and left to incubate for 10 minutes before addition 560 μL of 100% EtOH. RNA extraction was subsequently completed on-site according the manufacturer’s protocol for all sample types. To avoid contamination between samples due to the high number of virions, regular glove changes were conducted and parafilm was used to seal the gap between collection tubes and QIAamp Mini columns (Qiagen) during centrifugation. Batches always contained only primate or only human samples and a negative extraction control was processed with every batch. Human samples were linked to a record and clinical information such as date of onset of symptoms, date of sample collection, municipality, state of residence, age, sex, residence type and, when available, vaccine and travel history.

### Real-time quantitative PCR (sub-studies I to III)

YFV reverse transcription quantitative real-time PCR (RT-qPCR) was performed on 121 samples using the Superscript III Platinum One-Step qRT-PCR System (Invitrogen) on a StepOnePlus Real-Time PCR machine (Applied Biosystems). The conserved YFV 5’ non-coding region was targeted using the primers YFall15F (5’ to 3’: GCTAATTGAGGTGYATTGGTCTGC), YFall103R (5’ to 3’: CTGCTAATCGCTCAAMGAACG) and the probe YFall41 (5’ to 3’: FAM-ATCGAGTTGCTAGGCAATAAACAC-BHQ), based on the previously described Domingo’s assay (*48*). Thermocyler conditions consisted of reverse transcription at 45°C for 15 min, denaturation at 95 °C for 5 min, followed by 40 cycles of denaturation at 95 °C for 10s, and annealing and extension at 60°C for 40s. To check RNA isolation efficiency, we used RNase P as an endogenous positive control. Assays for RNase P used the primers RNaseP-F (5’ to 3’:AGATTTGGACCTGCGAGCG), RNaseP-R (GAGCGGCTGTCTCCACAAGT), and a probe (FAM-TTCTGACCTGAAGGCTCTGCGCG-BHQ1).

**Table S3.**
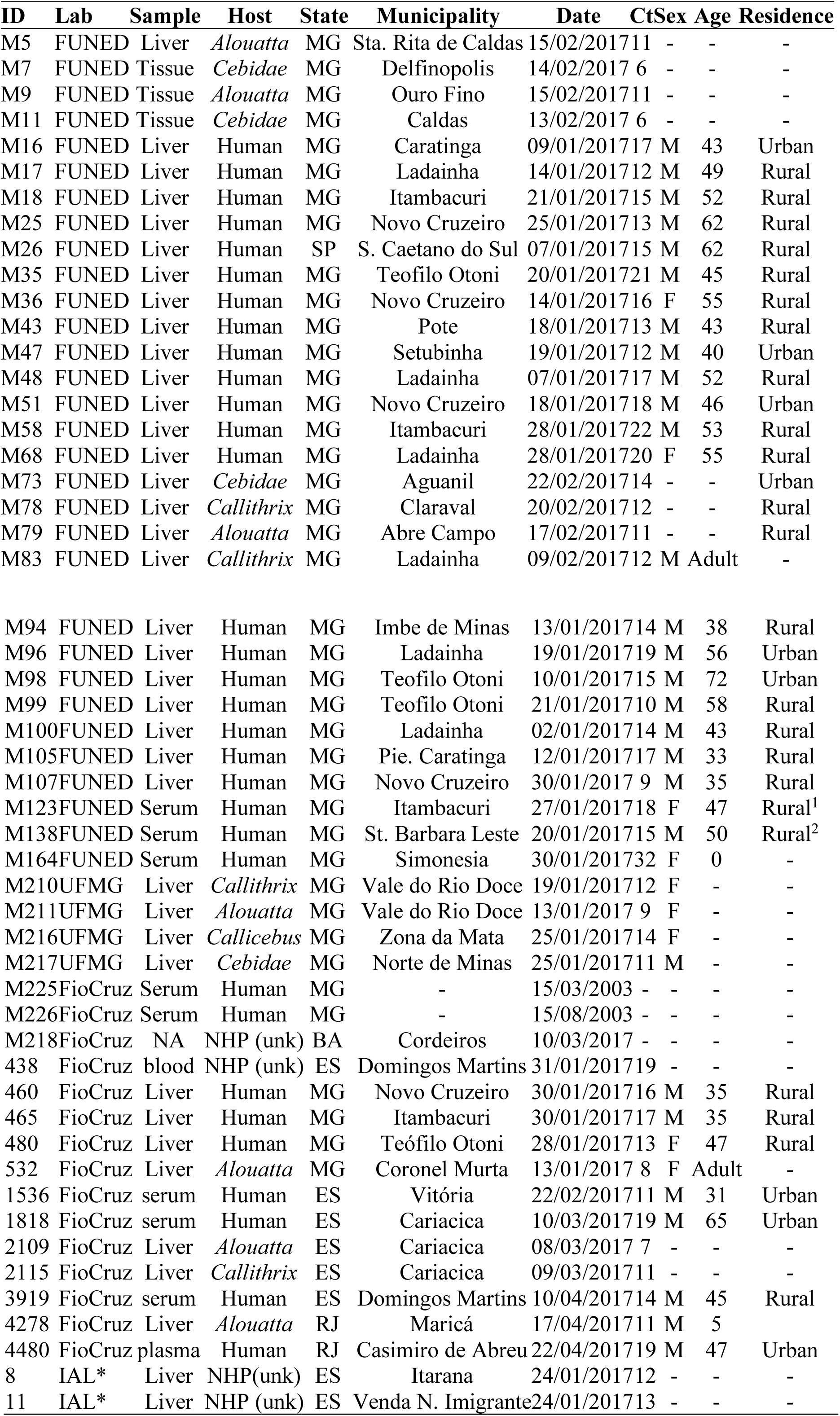
Epidemiological data associated with each isolate processed/sequenced in this study. ID=Project identifier; Lab=Laboratory where samples were processed/sequenced; Host=Host species; State: MG=Minas Gerais, BA=Bahia; ES=Espirito Santo; RJ=Rio de Janeiro; Date=Date of sample collection; Ct=RT-qPCR Cycle threshold value. “–”=not available. ^∗^*Alouatta spp.* suspected. ^1^=No date of vaccination available; patient was from São Paulo and was visiting Itambacuri (MG). ^2^ = Patient vaccinated in Jan 2017 and developed symptoms 3 days later.

### Validation of the sequencing primer scheme for MinION

Two candidate sequencing primer schemes were designed using Primal Scheme (http://primal.zibraproject.org) to amplify 500 bp or 1000 bp overlapping amplicons (*26*) of the complete genome of the YFV South American genotype 1, based on previous reports (*27*), with an overlap length of 75 bp between each neighbouring pair of primers. The scheme was validated at Public Heath England, UK. cDNA synthesis and multiplex PCR were conducted on RNA extracts from a cultured vaccine strain YFV 17D. PCR products were cleaned using 0.8x Ampure XP (Beckman Coulter) bead cleanups, quantified, and pooled. Libraries for the MinION were constructed using the ligation sequencing kit 1D (SQK-LSK108) and native barcoding kit (EXP-NBD103). The library was sequenced on an R9.4 flow cell (FLO-MIN106). Basecalled reads were aligned to a YFV reference genome using bwa (GenBank accession JF912190). Given that the regions overlap, alternate amplicons are amplified in two separate PCR reactions. These are pooled and barcoded together (in previous studies (*26*, *49*) these pools were barcoded separately, but this reduces the number of samples per flowcell by half). Mapping the reads to the reference genome showed the scheme provided good coverage across most of the coding-region of the genome. 95% of the genome had a depth of at least 379 reads, and 70% of the genome had a depth of at least 1941 reads. Both the 500bp and 1000bp PCRs with 40 cycles of PCR were tested in May 2017 at Minas Gerais (FUNED) on 7 samples of previously extracted RNA. Following PCR, quantitated dsDNA concentrations were higher for the 500 bp scheme than for the 1000 bp scheme, and therefore this scheme was chosen for all following assays (https://github.com/zibraproiect/zika-pipeline/tree/master/schemes).

### cDNA synthesis, library preparation and sequencing for MinION (sub-study I)

cDNA was reverse transcribed from viral RNA using the Protoscript II First Strand Sequencing kit (NEB) with random hexamer priming. Multiplex PCR was conducted using Q5 High Fidelity Hot-Start DNA Polymerase (New England Biolabs) and the 500bp sequencing primer scheme (**Table S4**). All samples were subjected to 32-40 cycles of PCR using the thermocycling conditions and reaction conditions described in Quick et al. (*26*). PCR products were purified using a 1x Ampure XP bead cleanup and concentrations were measured using a Qubit dsDNA High Sensitivity kit on a Qubit 3.0 fluorimeter (ThermoFisher). Library preparation for the ONT MinION was conducted using Ligation Sequencing 1D (SQK-LSK108) and Native Barcoding kit (EXP-NBD103) according to the manufacturer’s instructions, but with the changes detailed in (*26*). Amplified DNA and appropriate negative controls were sequenced in barcoded multiplexes of 6–12 samples per MinION run using FLO-MIN106 flow cells. Sequencing was performed without basecalling for 48 hours using MinKNOW. Consensus sequences for each barcoded sample were generated following previously published methods (*26*). Briefly, raw files were basecalled using Albacore, demultiplexed and trimmed using Porechop, and then mapped with *bwa* to a reference genome (GenBank Accession No. JF912190). Nanopolish variant calling was applied to the assembly to detect single nucleotide variants to the reference genome. Consensus sequences were generated; non-overlapped primer binding sites, and sites for which coverage was <20X were replaced with ambiguity code N. Sequencing statistics can be found in **table S5**.

**Table S4.**
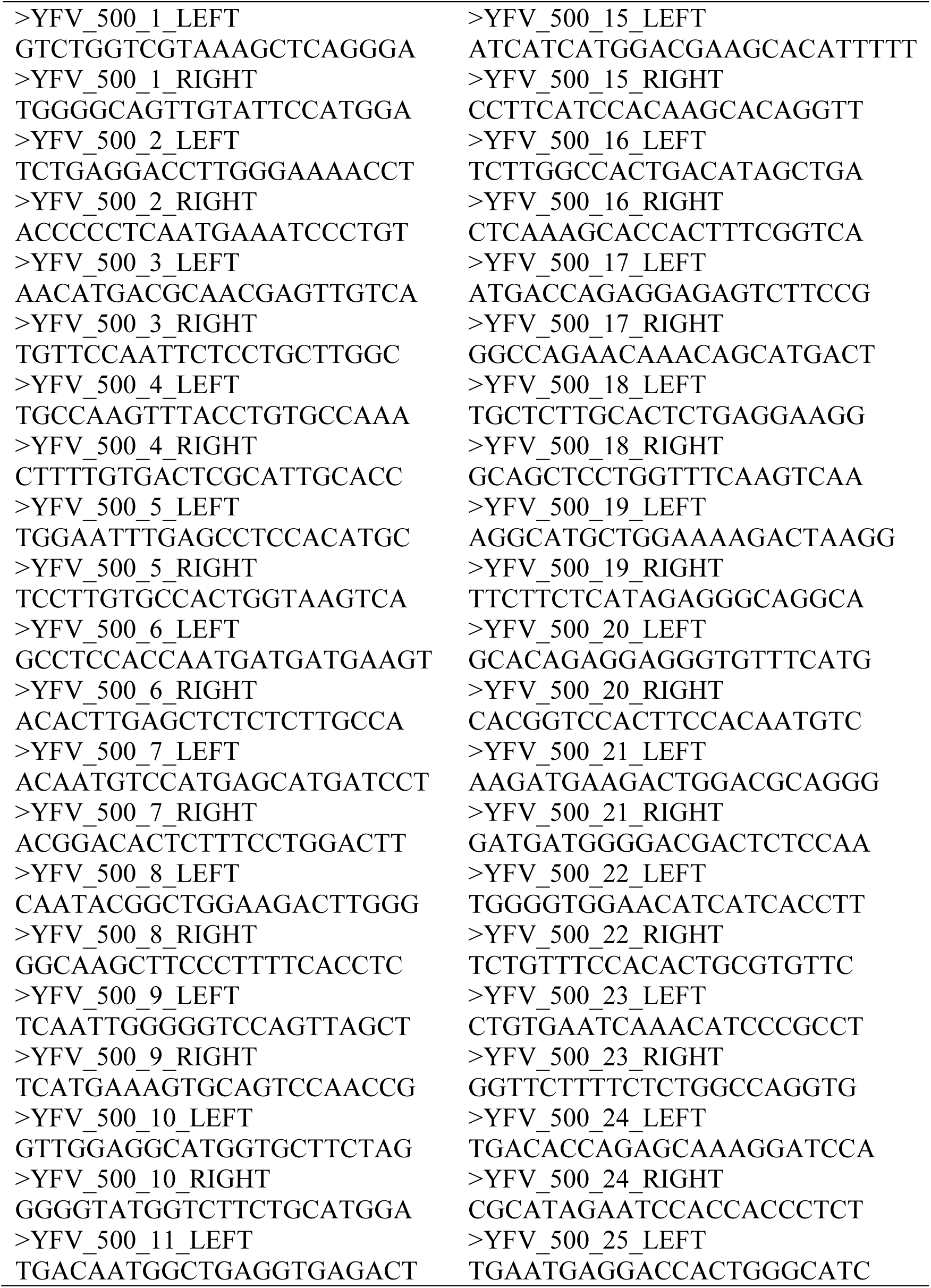
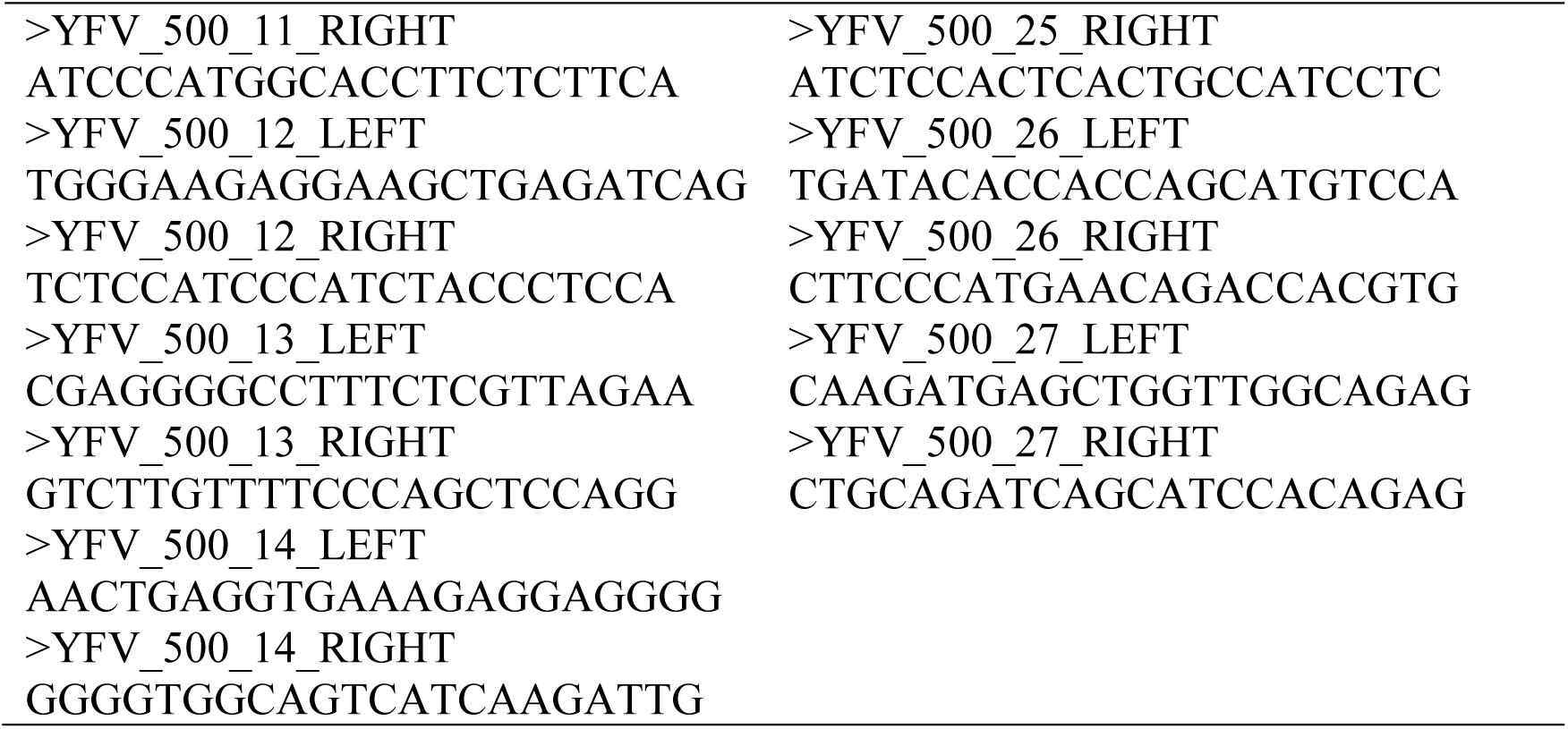
Primer sequences (*n*=*54*) for the YFV-500bp MinION sequencing scheme.

**Table S5.**
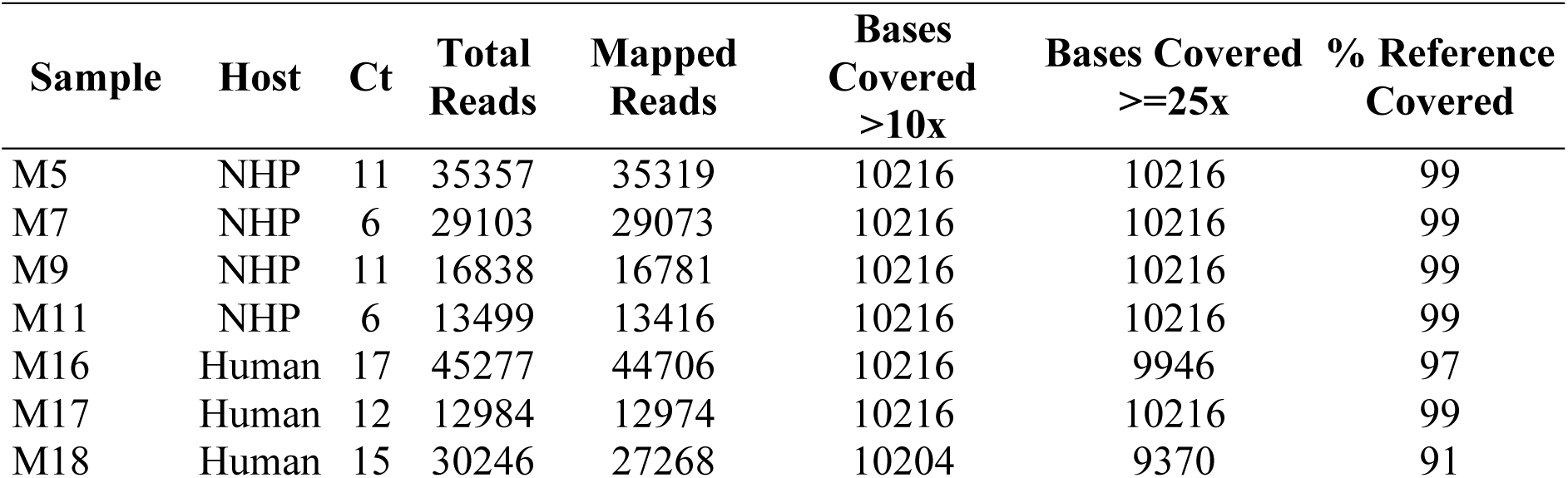
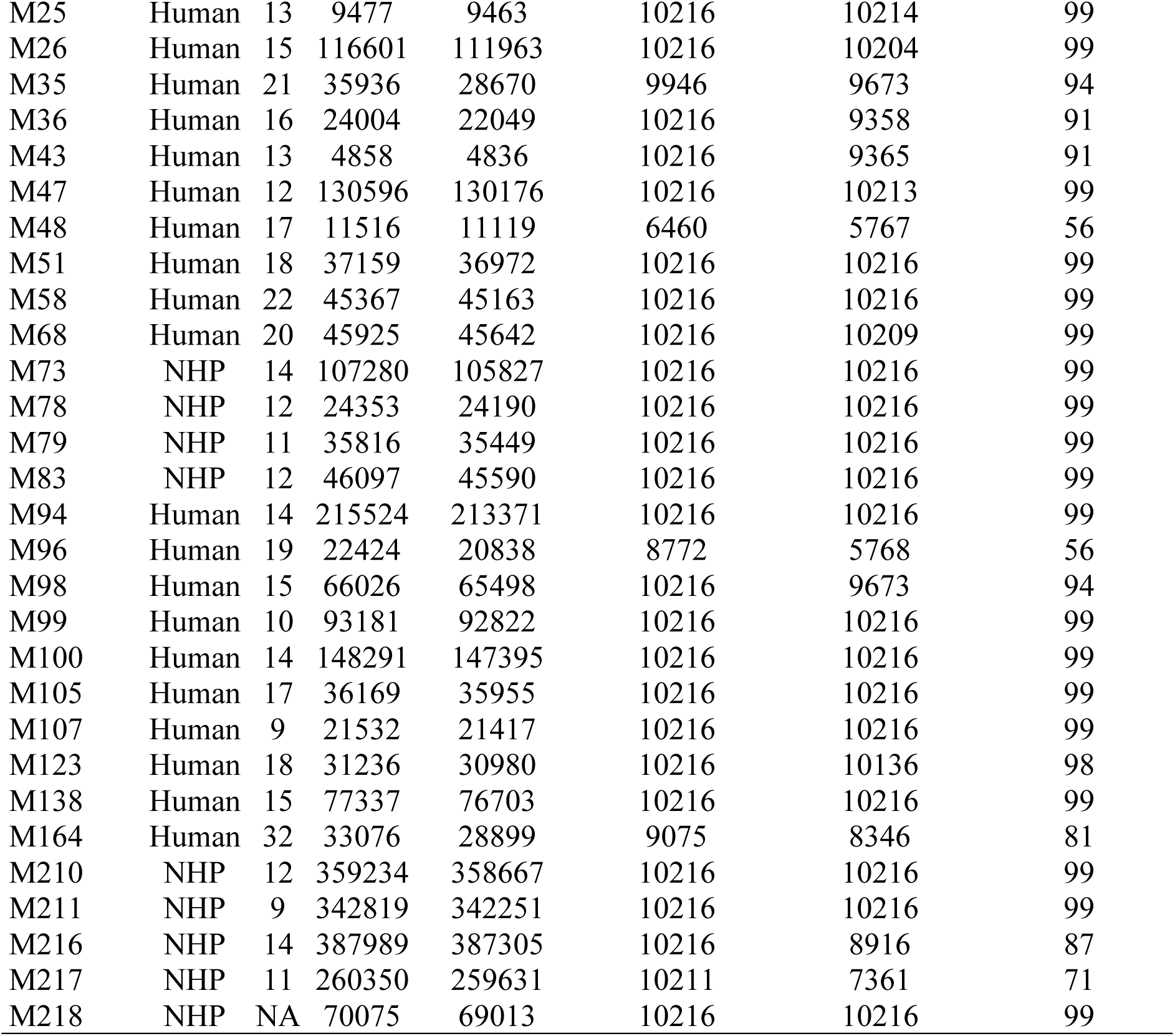
Statistics for the sequences generated using the MinION sequencer.

### cDNA synthesis and sequencing using Ion Torrent (sub-study II)

cDNA synthesis was executed with SuperScript III Reverse Transcriptase (ThermoFisher Scientific) and random hexamers. Subsequently, YFV genome amplification was performed using Platinum Taq High Fidelity DNA Polymerase (ThermoFisher Scientific). PCR products were analyzed by electrophoresis gel and purified using PureLink Genomic DNA spin columns (ThermoFisher Scientific). YFV amplicons were quantified using a Qubit dsDNA High Sensitivity kit on the Qubit Fluorometric 2.0 (ThermoFisher Scientific). Sequencing libraries were prepared using 100ng of PCR products with an Ion Plus Fragment Library Kit (ThermoFisher Scientific), according to the manufacturer’s instructions. For template amplification, emulsion PCR (emPCR) was performed using the Ion PGM Template OT2 kit and the Ion OneTouch 2 system (ThermoFisher Scientific). Ion Sphere particles (ISPs) were enriched using the Ion OneTouch ES (ThermoFisher Scientific). Enriched ISPs were sequenced using the Ion Torrent Personal Genome Machine Sequencer and the Ion PGM Hi-Q Sequencing kit (ThermoFisher Scientific), with the Ion 316 Chip. Data were collected for up to 8-9h. Reads were extracted, primer trimmed and mapped to a reference using Geneious R9 (9.1.7 version) (*50*). Briefly, primers were trimmed from each read (first 22 nt from 5’ end). Reads were extracted based on amplicon size and coverage normalization was performed. Consensus genome sequences were generated by reference mapping to GenBank accession JF912190.

### cDNA synthesis, library preparation and sequencing using Illumina (sub-study III)

Fourteen specimens were centrifuged at 20,000×g for 20 min and then filtered through a 0.45 μm filter (Merck Millipore, USA). The filtrates were treated with a mixture of nuclease enzymes to reduce background nucleic acids from the host cells and bacteria. RQ1 RNase-Free DNase (Promega Inc), DNase I (Zymo Research), Benzonase (Merck Millipore), RNase A (Zymo Research), RNase ONE (Promega Inc), Turbo DNase (Thermo Fisher) and 10X Turbo DNase buffer were added to the clarified supernatant and incubated at 37°C for 2h. Viral nucleic acids were extracted using a Maxwell 16 automated extractor (Promega Inc). Viral cDNA synthesis from extracted viral RNA/DNA was performed by using 50 pmol of a dodecamer of random primer in a reverse transcription reaction with AMV Reverse Transcriptase (Promega Inc) and RNasin Ribonuclease Inhibitor (Promega Inc). The 2nd strand cDNA synthesis was performed using DNA Polymerase I Large (Klenow) Fragment (Promega Inc), followed by the use of a Nextera XT Sample Preparation Kit (Illumina Inc) to construct a DNA library with each sample identifiable using dual barcodes. For size selection, we used a Pippin Prep (Sage Science Inc) to select a 400 bp insert (range 200-600 bp). The library was deep-sequenced using the MiSeq Illumina platform with 2 × 300 bp paired ends. Paired-end reads of 2×300 bp generated by MiSeq were demultiplexed using the vendor software from Illumina. Demultiplexed Illumina reads were mapped on the JF912190 reference genome using bwa-mem program (*51*). The genome analysis toolkit (*52*) was used to perform variant calling and generate consensus sequences with a 3x minimum read depth coverage.

### Automated phylogenetic typing tool

We developed an tool that automatically classifies and accurately annotates YFV genome sequences, which is publicly available at http://bioafrica2.mrc.ac.za/rega-genotype/typingtool/yellowfevervirus/.

To build this YFV typing tool, we prepared two reference datasets that include publicly available sequences, one with whole-genomes (*n*=34, length=10,235 bp) and another with envelope gene sequences (*n*=34, length=1,443 bp). The accession numbers for each reference sequence of each genotype are as follows are as follows; for South American genotype 1: JF912190, JF912187, JF912188, JF912189, JF912180, JF912182, JF912185, JF912179, JF912184, JF912183, JF912186; for South American genotype 2: TVP17388, JF912181; for the West African genotype: AF094612, JX898871, JX898872, AY640589, JX898875, JX898874, JX898873, AY572535, AY603338, JX898868, JX898870, JX898876, JX898878, JX898880, X898877, JX898869, YFU54798; and for the East African genotype: AY968064, AY968065, DQ235229, JN620362. To validate the reference datasets, phylogenetic trees were constructed using maximum likelihood (ML) with a general time-reversible model and among-site rate variation modeled using a discretized gamma distribution (GTR + Γ_4_), which was inferred as the best-fitting nucleotide substitution model in jModelTest (*53*). Trees were estimated using RAxML v8 (*54*) with 100 bootstrap replicates. All genotype clades are supported by bootstrap values of 100%, with the exception of the West-African genotype in the *env* tree, which is supported by a bootstrap score of 99%.

Classification of query sequences using the YFV subtyping tool involves two steps. The first step identifies the virus species using the basic local alignment search tool (*55*) that searches the RefSeq NCBI Reference sequence database that contains 7952 viruses reference genomes (*56*). The virus species is identified if the alignment score >400, which is the sum of identities minus gaps and mismatches. In addition, the tool also creates a codon alignment and identifies polymorphic sites and genetic diversity in the alignment, and aligns the query sequence to the NC_002031 curated reference sequence (*57*).

The second step involves the reconstruction of a phylogenetic tree with a reference dataset using neighbour-joining (**fig. S4**). Statistical support for phylogenetic clustering of the query strain with the pre-defined reference genotypes using 1,000 bootstrap replicates. A query sequence is assigned to a particular genotype if clustering is supported by a bootstrap score >70%. The YFV typing tool accepts up to 2,000 sequences per submission and analyses each of sequence independently. At the end of the analysis, a phylogenetic tree is created that displays all query sequences and the reference dataset. A formatted report, estimated phylogenetic tree, and alignments can all be downloaded in multiple formats by the user.

**Fig. S4.**
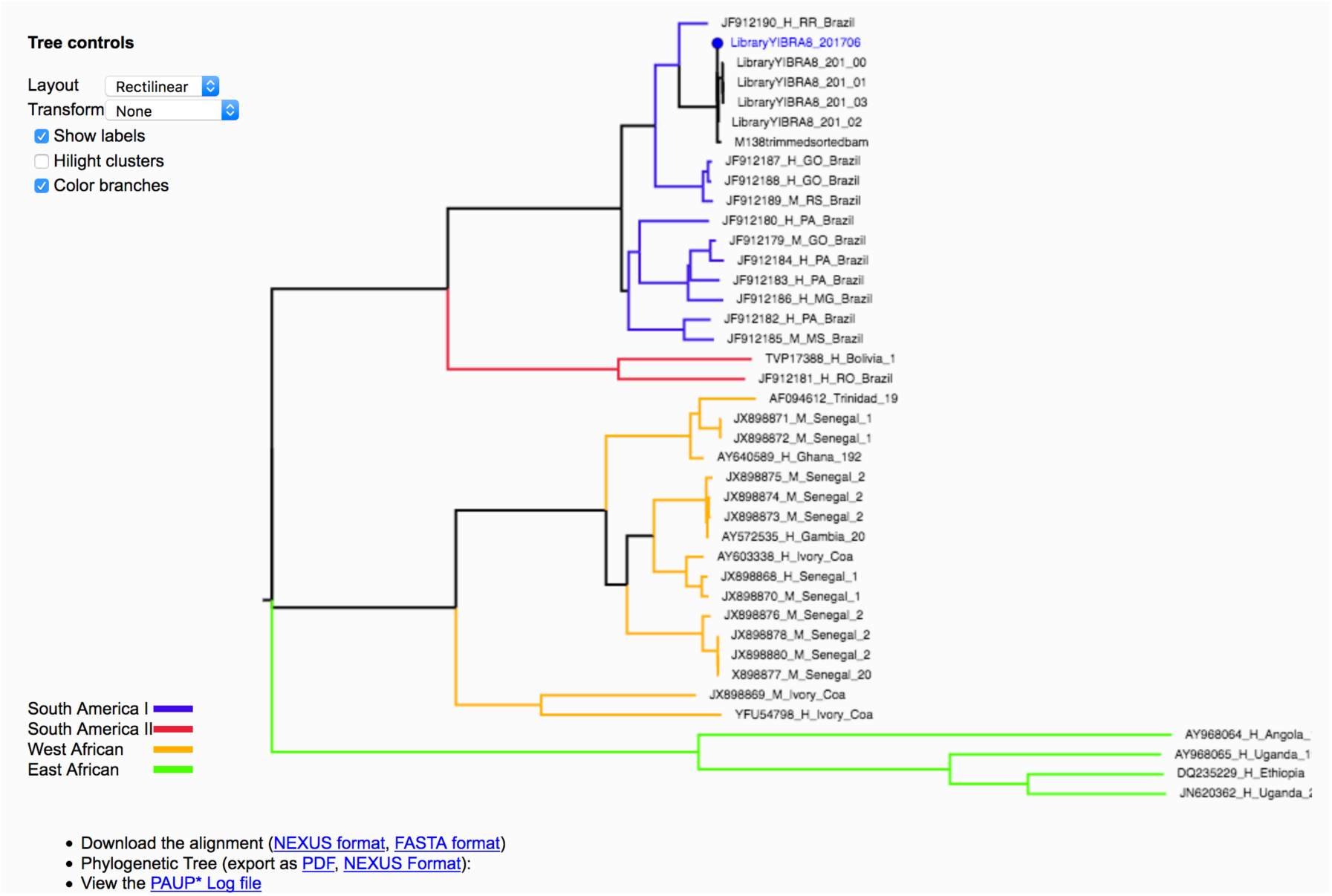
Illustration of the output of the online YFV classification tool. The figure shows the ML phylogeny of 6 target sequences analyzed by the tool. The output also provides a link to genome coverage and a more detailed report. The reference dataset is colored according to genotype.

### Curation of whole-genome sequence datasets

We screened the GenBank database for published complete YFV genome sequences sampled worldwide using an in-house shell script. We retrieved publicly available data from a total of 14 countries across several regions: Caribbean (Trinidad), East Africa (Ethiopia, Uganda and Sudan), Central Africa (Angola), East Asia (China), West Africa (Senegal, Ivory Coast, Ghana, Nigeria and Guinea-Bissau), and South America (Bolivia and Brazil). These genome represent viruses sampled over the last 90 years, from 1927 to 2017. Sequences were aligned using MAFFT v.7 (*58*). Maximum likelihood phylogenetic trees were estimated using RAxML (*54*) under a GTR + Γ_4_ nucleotide substitution model, as described previously. Root-to-tip regressions of sequence sampling date against genetic divergence were conducted (**fig. S5**) to identify and remove 7 (sub-study III) potential contaminants or mislabelled sequences (*59*). ML trees and root-to-tip analyses were performed for several datasets (**fig. S5**). All alignments were screened for recombination using the Phi-test available in SplitsTree v.4 (*60*); the null hypothesis of the absence of recombination could not be rejected (P<0.05) and lack of recombination was confirmed using the RDP4 package (*61*). The *outbreak dataset* comprises 52 genome isolates, 50 of which were generated by this study and 2 of which were published in (*27*) (clade c in **figure S5**; see also **table S3**).

**Fig. S5.**
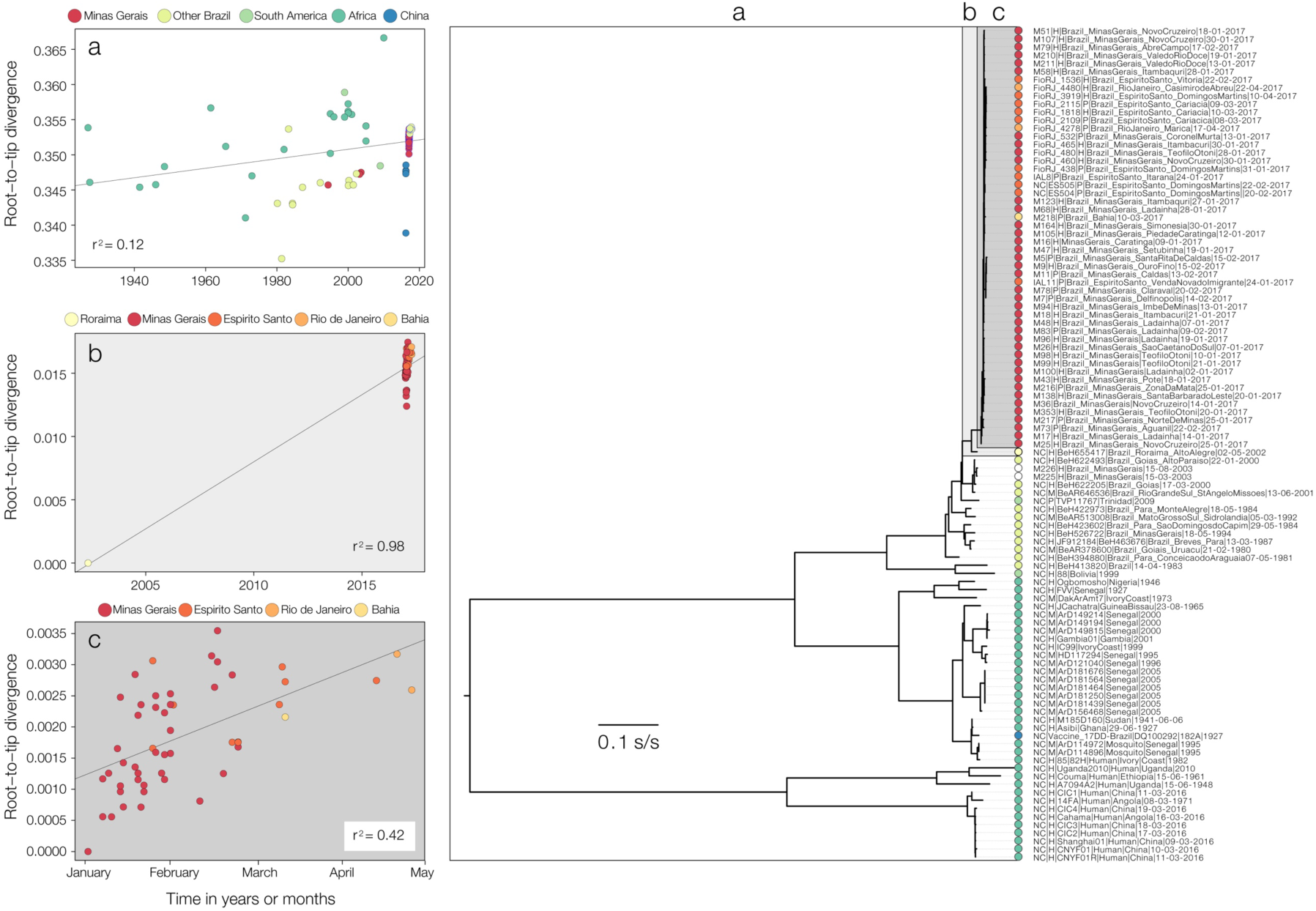
Temporal signal of YFV sequence datasets. Panels on the left show the correlations between sampling dates and genetic distances from the tips to the root of maximum likelihood phylogenies estimated for clades a, b and c (taxa belonging to each clade are indicated by the labeled boxes in the phylogeny to the right side). Phylogenies were estimated using RAxML (*54*) under a GTR+Γ_4_ nucleotide substitution model and correlations generated using TempEst (*59*). Tips are coloured according to administrative unit of sample collection, and taxa names are indicated on the right hand side of the phylogeny (P=non-human primate; H=Human, M=mosquitoes). Taxa names include strain name, host species, location and date of isolate collection.

### Bayesian skygrid with epidemiological covariates

Time-scaled phylogenetic trees were reconstructed using the Bayesian phylogenetic inference framework available in BEAST v1.8.4 (*62*). We used a fully probabilistic approach that combined sequence substitution over an unknown tree, calibrated to a real time scale using a molecular clock model. We used the HKY + Γ_4_ nucleotide substitution model and a relaxed molecular clock model, with an underlying lognormal distribution of branch rates (*63*).

For the molecular clock model, we assumed that the outbreak clade exhibited a different clock rate to ancestral paraphyletic lineages, as observed in previous epidemics (*64*) and therefore we used a fixed local clock model (27) on clade B (comprising 66 South American genotype 1 whole genomes; **fig. S5**). We also computed a Bayesian skygrid with covariates model using the outbreak clade A sequences alone (see **fig. S5**), for which we specified 36 grids (i.e. the approximate number of epidemiological weeks spanned by the duration of the phylogeny). Further, we ran a Bayesian skygrid-based generalized linear model (*31*) with a streamlined prior specification in which effective population size through time is associated with a single covariate, chosen probabilistically from a set of possible covariates, while also accounting for phylogenetic uncertainty. In this analysis we investigated the following set of 3 covariates: i) log-linear YFV(H) case counts (dataset A), ii) log-linear YFV(NHP) case counts (dataset B) and iii) log-linear CHIKV human case counts (dataset C). Specifically, for each grid point (epidemiological week) we include the log-transformed and standardized number of cases as described in **Section 1**. The association of each particular covariate with the effective population size dynamics of the outbreak is summarised by an inclusion probability. Distributions of the outbreak TMRCA obtained without and with covariates are shown in **Fig. 3c** (distributions b and c, respectively). Further, a comparison of the TMRCA estimates with and without YFV case counts can be found in **fig. S6**.

**Fig. S6.**
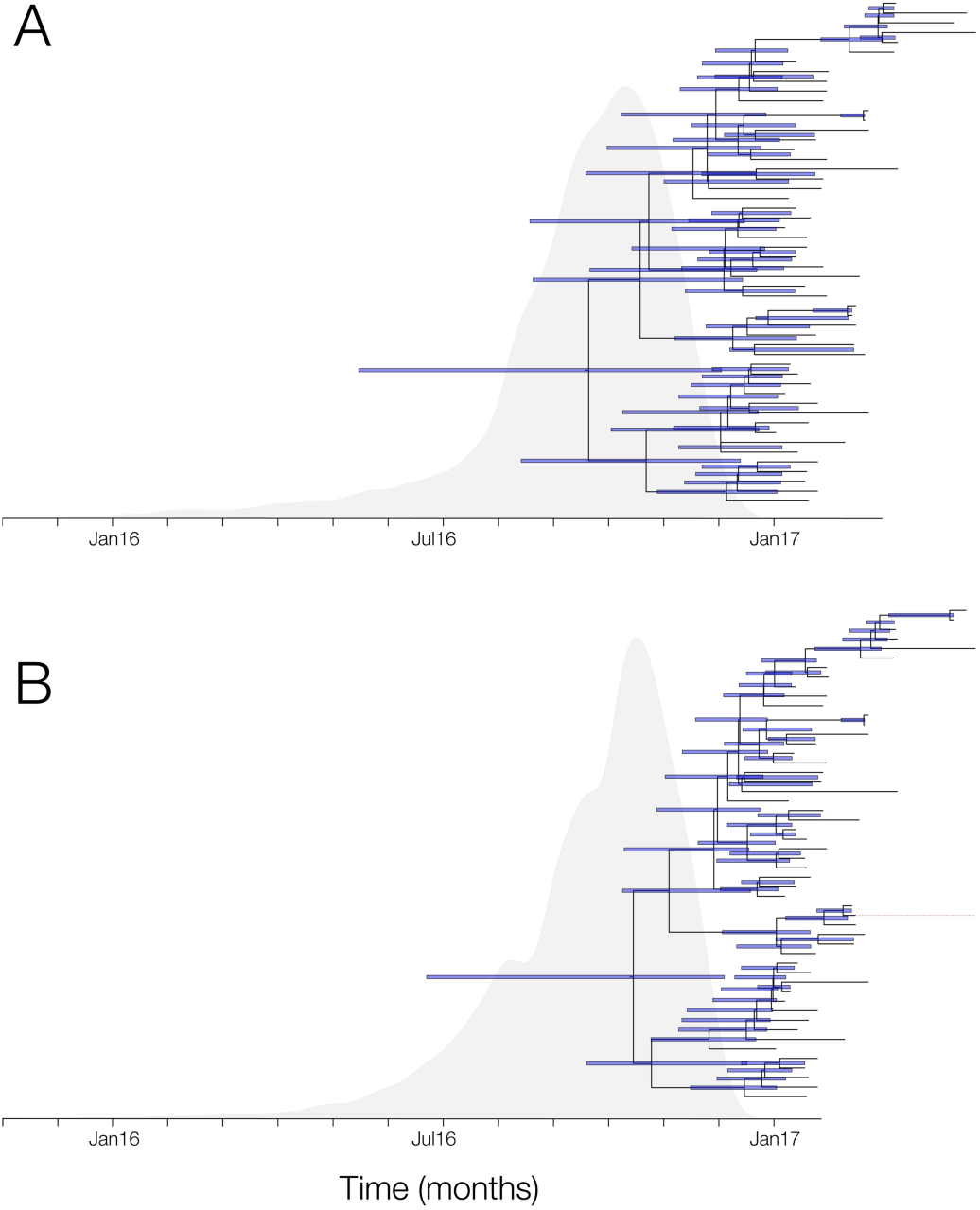
Combining virus phylogeny and epidemic time series. Maximum clade credibility trees (MCCs) generated in BEAST are shown together with the corresponding posterior distribution of the time of most recent common ancestor (TMRCA; grey) of the outbreak clade. Panel (A) shows the results obtained using the standard skygrid model whilst (B) shows the results obtained using skygrid model with covariates (B). Addition of the epidemiological time series data reduces the statistical uncertainty of the estimated TMRCA parameter by 22%. Purple bars indicate uncertainty in estimates of divergence times of internal nodes in each tree.

### Structured coalescent analyses

Viral lineage transitions among hosts were inferred using a structured coalescent model, as implemented in the MultiTypeTree v6.3.0 package (*32*) for BEAST v2.4.7 (*65*). The analysis was performed on the “*outbreak dataset*” only (i.e. clade c indicated in **Fig. S5**). The structured coalescent model also estimates time-scaled phylogenetic trees and state transition histories. It assumes a constant effective population size for each deme (i.e. human vs nonhuman host states, in this study) and asymmetric transition rates between demes. As in the other analyses above, we used an uncorrelated relaxed molecular clock model with a lognormal distribution prior on the branch rate parameters (*63*) and a HKY+Γ_4_ nucleotide substitution model. Default priors were used for the nucleotide substitution model. A lognormal prior was placed on the molecular clock rate parameter, with mean equal to 0.001 susbtitutions per site per year (in real space) and standard deviation set to 1. An exponential prior with mean 1 was used for the effective population sizes of demes and transition rates between demes (=host species states). To ensure that the phylogenetic timescale is well informed we placed a normally distributed prior with mean 0.6306 years before the present (and standard deviation 0.11) on the time of the most recent common ancestor (TMRCA) of the tree. This TMRCA prior covers the 95% HPD interval of the TMRCA inferred by the skygrid model with the best-fitting covariate (i.e. posterior distribution c in **Fig. 3c**). When estimating transition rates between host states, two independent runs of 200 million steps were computed, sampling parameters every 20,000 steps. The two chains were combined with LogCombiner, discarding 10% of each chain as burn-in and subsampling only half of the remaining states. Tracer v1.6.0 (http://tree.bio.ed.ac.uk/software/tracer/) was used to check the MCMC analysis for convergence. A maximum clade credibility tree with annotated branches was then generated in TreeAnnotator (**Fig. 4A**; the same tree with detailed taxa information is shown in **fig. S7**). To recover the number host-switching events through time we counted the number of transitions between demes (host states) across monthly intervals for each tree in the posterior set of structured coalescent trees (migration histories). This count and its 95% HPD interval are shown in **fig. 4C**.

**Fig. S7.**
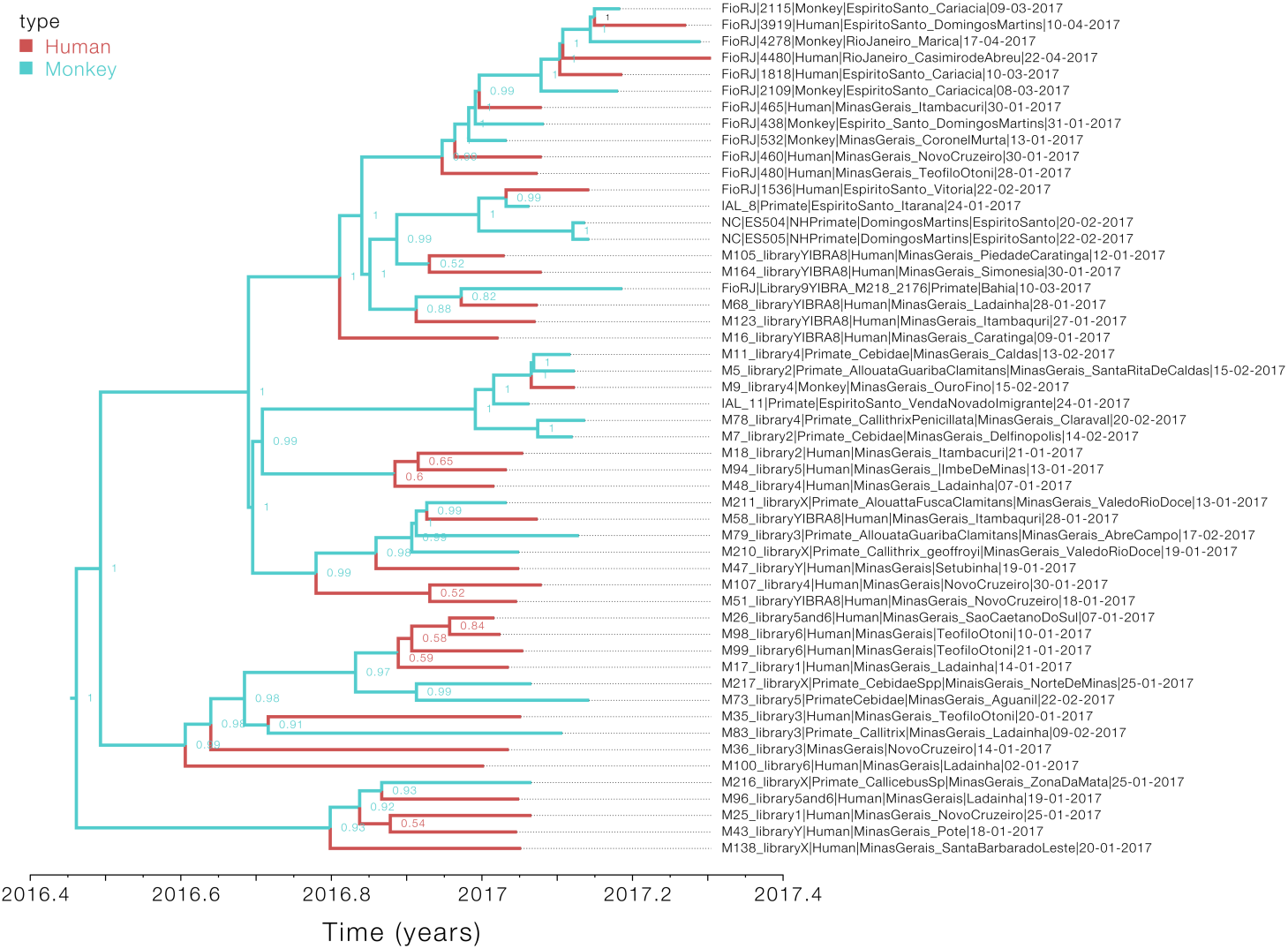
Typed maximum clade credibility tree (corresponding to Fig. 4A). The node labels indicate the posterior probability of the most likely host state for each internal node, inferred by the MultiTypeTree package (*32*). This representation does not include information on individual transition events between host states.

To test sensitivity to the TMRCA prior used, the analysis was repeated (i) without a prior on the TMRCA and (ii) using a normally distributed prior with mean 0.837 years before the present and standard deviation 0.25, which corresponds to the TMRCA inferred under a standard skygrid model (i.e. without covariates; see **Fig. 3C**). The inferred posterior distributions of the transition rates between human and NHP host states are shown in **fig. S8**, where it can be seen that the TMRCA prior does not significantly affect the estimate transition rate dynamics. We also verified that this is the case for the migration histories (data not shown). In addition, the rate of host-transition events from NHP to human - our key result - always clearly deviates from the prior, whereas the reverse rate (from human to NHP) recovers the prior. To further test the robustness of the estimated transition rates and the number of host-switching events through time, all analyses were repeated using a lognormal prior (with mean 0 and standard deviation 4) on the deme effective population sizes and on the transition rates between demes. This resulted in estimating slightly higher rates and thus inferring more host-transition events. Nonetheless, the HPD intervals of host transition rates under different priors largely overlap and the overall pattern was similar (not shown).

**Fig. S8.**
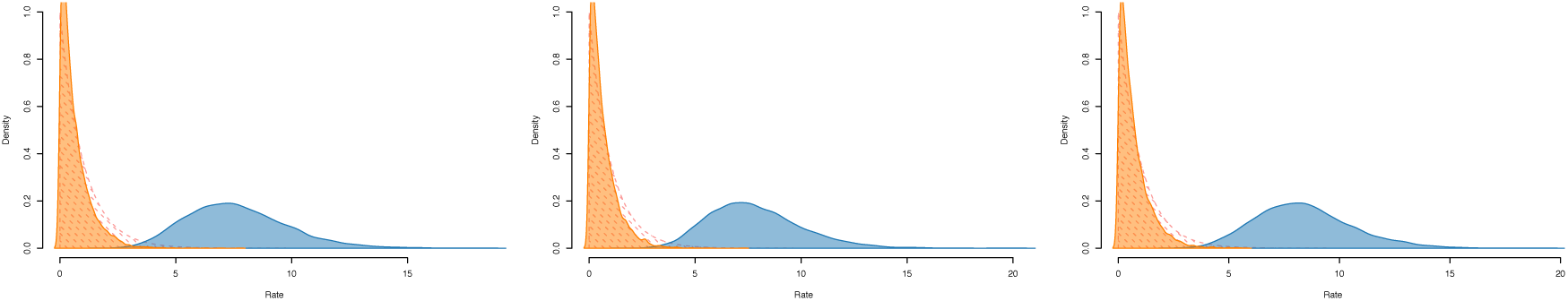
Estimated posterior distributions of the host state transition rates of the structured coalescent model under different priors. From left to right, (i) using no prior on the TMRCA, (ii) using a normal distribution with mean 0.631 years before the present and standard deviation of 0.11, and (ii) using a normal distribution with mean 0.837 years before the present and a standard deviation of 0.25. The prior distribution used for the migration rates is shaded with dashed red lines. Orange=human to non-human primates spillovers, blue=NHP to humans spillovers.

### Phylogeographic inference in continuous space

Bayesian continuous phylogeographic analyses were performed on the “*outbreak dataset*” only (i.e. clade c in **Fig. S5**) using the skygrid with covariates as the coalescent tree prior (*31*). We first inferred the best fitting continuous diffusion process by performing (log) marginal likelihood estimation using generalized stepping-stone sampling (*66*) on a range of relaxed random walk models, as well as the time-homogeneous Brownian motion process (**table S6**). Details of the stepping stone sampling approach were as follows: after an initial posterior exploration of 10 million iterations, we collected 1,000 samples from each of the 51 power distributions, distributed according to a Beta(0.3,1.0) distribution and sampling at every 1000^th^ iteration. The log marginal likelihood estimates were highly consistent between independent runs in BEAST1.8.4 (*67*).

**Table S6.** Generalized stepping-stone (GSS) sampling for each of the continuous diffusion models. Models are ordered according to their Bayes Factor (BF) score, calculated against the Brownian diffusion model (homogeneous diffusion).

| **Model** | **GSS** | **Bayes Factor** |
| --- | --- | --- |
| Homogeneous | -16137.39 | 0.0 |
| Lognormal | -16082.79 | 54.60 |
| Gamma | -16082.01 | 55.38 |
| **Cauchy** | **-16075.22** | **62.17** |

All the relaxed random walk models strongly outperformed the time-homogenous Brownian diffusion model. A model with Cauchy distributed diffusion rate variation among branches yielded the highest Bayes Factor (BF) against the time-homogenous diffusion model, indicating among-branch heterogeneity in branch velocity. The Cauchy model is strongly preferred among all the relaxed random walk models (**table S6**).

The Cauchy-distributed phylogeographic model selected above was then used to characterise the outbreak’s spatio-temporal epidemic history (*34*). Posterior distributions under the Cauchy models were obtained using Markov chain Monte Carlo (MCMC) sampling as implemented in BEAST v 1.8.4 (*62*). The BEAGLE library v2.1.2 was used to accelerate computation (*68*). MCMC chains were run in triplicate for 250 million generations, sampling every 50,000 steps. MCMC performance was inspected for convergence and for sufficient sampling using Tracer v.1.6.

To summarise virus diffusion over time and space, 1,000 post-burn-in phylogenies sampled at regular intervals from the posterior distribution were obtained. The branches of these phylogenies were extracted as vectors, each having start and end spatial coordinates, and start and end dates (i.e. branch duration) in decimal units (*36*). The R package “seraphim” was used to estimate statistics of spatial dissemination, such as dispersal velocity, diffusion coefficients, and evolution of the maximal wavefront distance from epidemic origin (*68*, *69*), as well as generating monthly graphical representations of the inferred spatio-temporal spread process (**Movie S1**) using the “spreadGraphic” function (*70*).

### Data availability

Epidemiological case counts, genome alignments, BEAST XML files, and code used in this study can be downloaded from the ZiBRA Github website (https://github.com/zibraproject; MinION sequencing protocols can be found at https://github.com/zibraproject/zika-pipeline/tree/master/schemes). Genome sequences generated in this study are publicly available in GenBank database under the accession numbers: MH018064-MH018115.

